# How repair-or-dispose decisions under stress can initiate disease progression

**DOI:** 10.1101/828053

**Authors:** Andreas Nold, Danylo Batulin, Katharina Birkner, Stefan Bittner, Tatjana Tchumatchenko

**Affiliations:** Theory of Neural Dynamics, Max Planck Institute for Brain Research, Frankfurt, Germany; Department of Neurology, Focus Program Translational Neuroscience (FTN), University Medical Center of the Johannes Gutenberg University Mainz, Mainz, Germany

## Abstract

Glia, the helper cells of the brain, are essential in maintaining neural resilience across time and varying challenges: By reacting to changes in neuronal health glia carefully balance repair or disposal of injured neurons to prevent further tissue damage. Malfunction of these interactions is implicated in many neurodegenerative diseases. Reductionist models with a minimal number of parameters provide the opportunity to gain insight into biological functions and inform experimental designs. We introduce such a model that mimics long-term implications of repair-or-dispose decisions. Depending on the functionality of the decision-making process, the model assumes four distinct tissue states: healthy, challenged, primed tissue at risk of acute damage propagation, and chronic neurodegeneration. These states of the model correspond to the progression stages observed in the most common neurodegenerative conditions. The underlying mechanisms are in agreement with experimental observations of glia-neuron crosstalk and reproduce a homeostatic balance between repairing and damage-inducing reactions. The model suggests that the onset of neurodegeneration results from a tug-of-war between two conflicting goals: short-term resilience to stressors vs long-term prevention of tissue damage.

## Introduction

Throughout our lifetime the brain faces the challenge of keeping neuronal networks functional under widely varying changing conditions. This requires repairing injured neurons, or if their damage is irreversible, disposing of them in order to maintain global tissue health. This decision is made largely by glia-neuron crosstalk in the brain and needs to be critically balanced because neurons are postmitotic cells and stem-cell niches have only a limited potential for regeneration. One of long-standing challenges in clinical neuroscience is to understand how the brain upholds its amazing resilience capacity, and how glia-neuron interactions contribute to it.

As a diverse set of cells, glial cells consist of astrocytes, microglia and oligodendrocytes among others, each playing a vital role in protecting neurons from harm, supplying nutrients, and establishing an environment where neurons can thrive. For example, astrocytes respond to threats such as external trauma or inflammation by transitioning to a ‘reactive’ pro-inflammatory phenotype. Microglia constantly surveil their local microenvironment, clear the tissue of pathogens and phagocyte synapses or dead cell material. Oligodendrocytes help maintain axonal health and defend against the threatening factors to axonal integrity by eliciting an increase of precursor cell differentiation and increasing remyelination.

Importantly, neurons communicate directly with glia to signal their health state and either suppress or increase glial activity. The endogenous glia-neuron feedback-loop normally responds to perturbations in a way that a lesion is contained and neuronal network function is maintained. Malfunction of this feedback can be triggered by a variety of causes ranging from trauma, inflammation, stroke, break-down of the blood brain barrier to aging, ultimately to neurodegeneration and functional deficits. Upon intrinsic and external stressors, the severity of the neuronal loss and how quickly it spreads depends significantly on the functionality of the glia-neuron feedback loop – suggesting a critical role for this communication in maintaining brain resilience.

One crucial aspect for the functionality of glia-neuron feedback is the balance between repair and dispose-inducing tissue reactions. In the healthy tissue, both modes are tightly controlled by checkpoint mechanisms and a web of complex cellular interactions. We consider the net effect of this complex array of interactions: The repair or disposal of damaged neurons. We put forward an abstract model for this process, and study the implications that a dysregulation of the repair-or-dispose decision process has for disease progression. By following a reductionist approach, we aim to uncover basic mechanistic principles which could be responsible for disease progression. But, how viable is it to model a process of enormous complexity such as the glia-neuron crosstalk in the context of highly complex and diverse neurodegenerative diseases, through a reductionist mathematical model?

In the last couple of decades, several modeling approaches have been put forward to better understand processes underlying neurodegeneration and neural tissue resilience [1, 2, 3]. The modeling approaches vary widely and address aspects from nervous system energy metabolism [4, 5], protein degradation [6, 7], blood-brain barrier transport [8, 9] to tauopathy [10, 11]. For example, studies that focus on the role of a particular signaling pathway typically aim to capture a specific biochemical process as accurately as possible [12, 10, 13, 14]. Typically, the mathematical model consists of a set of biochemical reactions describing e.g. the interplay between protein synthesis, folding, refolding, degradation, and aggregation [14]. However, the detailed model design requires strong assumptions about the conditions in which it is valid, and often hinges on extensive use of in vitro data for parameter estimation and model verification. This limits the explanatory and predictive power for the overall disease progression. Computational models that target processes by multiple agents and/or on several timescales, require either higher levels of abstraction.

One such approach is followed by data-driven models. Such models can, for example, predict the mortality risk for traumatic brain injury patients based on clinical variables such as age, gender, and discretely monitored inflammatory biomarkers [15, 16]. However, the data-driven design principle often does not allow to infer a mechanistic understanding of underlying disease processes. To achieve this, abstract models are used. Abstract mechanistic models address a broader aspect of disease progression at the expense of detail by model reduction. In contrast to data-driven models, which infer quantitative predictions based on past observations, the primary goal of mechanistic models is to drive conceptual insights into disease processes and therefore generate hypotheses. For example, a neuronal mass model of partial epileptic seizures [17] models transitions between healthy nervous system activity and seizure-like events using only five state variables, and is used in basic research on epilepsy [18, 19] and in clinical studies [20, 21]. In the field of neurological diseases, diffusion models [22] and epidemic spread models for misfolded protein dynamics [23, 24] have linked the spread of toxic proteins in 3D brain structures with spatio-temporal patterns of pathology development. The main challenge for the development of abstract models is to determine an optimal abstraction level by the choice of state variables, which follows both existing evidence and allows for model verification using available experimental techniques. We believe that the understanding of glia-neuron crosstalk and neurodegenerative disease processes has advanced enough to put forward a mechanistic top-level model of the implications of repair-or-dispose decisions in glia-neuron crosstalk for long-term disease progression.

We adopt a reductionist computational model to understand how such a self-cleaning mechanism might work and fail (see Fig. S1). The model includes two coarse grained state variables of an assembly of cells. For the tissue to be functional, the number of eliminated cells needs to be limited. However, external stressors can induce local cell damage. If untreated, these cells then damage connected cells, therefore compromising functionality and stability of the cell assembly. The system therefore includes a self-cleaning mechanism by which damaged cells are disposed of. Model computations based on these core assumptions reveal four distinct states for the assembly: healthy, challenged, primed and chronic. Importantly, they show that with rising baseline damage, the self-cleaning mechanism needs to satisfy two conflicting goals: It needs to recognize and remove damaged cells and at the same time avoid collateral damage. The result of this tug-of-war between long- and short-term resilience determines how long the cell assembly can be maintained. The manuscript is structured as follows: We set up the model and present its functional modes in the Results section. We then discuss analogies to cellular processes and to experimental and clinical findings for neurodegenerative disease courses. Finally, we suggest a mechanistic explanation for the disease progression of Alzheimer’s disease and Multiple Sclerosis.

## Results

### Model

Similar to the spread of a virus from human to human in a pandemic, there are diseases in which damage propagates from cell to cell. For instance, in several neurodegenerative diseases, neurons can induce damage-inducing processes via synapses (see eg. Ref. [24, 25], and Discussion for further disease-specific processes). Such cells, which spread damage to connected cells, will be referred to as ‘seeds’. If not contained, then widespread tissue damage can initiate from a single seed. First, this seed increases damage in connected cells. Then, once these connected cells exceed a damage threshold, they become seeds themselves, and also start propagating damage. Because every cell spreads damage to many other cells, this process accelerates exponentially. We are interested in the following scenario: First, we consider cases where damage spread from one cell to another cell is slow, in the range of weeks to years, but tissue maintenance is fast, in the timescale of hours to days. Second, we are interested in capturing the point of ‘ignition’, when the first seed successfully transmits damage, as accurately as possible by resolving the state of individual cells.

To formalize the spread in this scenario, each cell (with index *n*) is attributed one variable *z*_*n*_ > 0. This variable represents the ‘baseline damage’ of the cell. We define the following three states based on *z*: If *z* < *z*_cliff_, the cell does not spread damage. If *z*_cliff_ < *z* < 1, it becomes a seed and spreads damage to connected cells. Once *z* = 1, the cell dies. The lower *z*_cliff_ is, the higher is the tissue susceptibility for slow damage spread. Here, for simplicity we consider the intermediate case *z*_cliff_ = 0.5. The following equation formalizes the core process of such a spreading process:

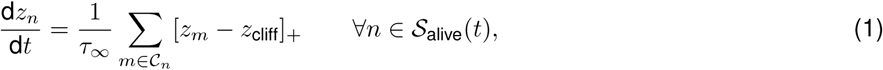

where *τ*_∞_ is the duration of cell-to-cell transmission and [*z*]_+_ = *z* for *z* > 0 and zero otherwise. Each cell ‘receives’ damage from *M* other cells. These potential damage-transmitters are randomly selected for each cell, and are defined by the set 𝒞 _*n*_. Once a cell dies, it is excluded from 𝒞 _*n*_ and can no longer transmit damage. The process of damage spread is initiated by externally driven irreversible ‘seeding events’. For example, this could be an abnormal tau accumulation in Alzheimer’s disease (see Discussion for details). We assume that these seeding events are rare and sparse, but that they will happen to some cells in the tissue. If left unchecked, they initiate devastating, exponentially increasing damage. Fig. 1 shows the damage spread, starting from a single external seeding event. First, the seed induces damage to its connected cells, which then in their turn spread damage. This cascade leads first to an exponential increase in the number of seeds, and then an exponential increase of cell deaths. How can this undesirable result be avoided?

**Figure 1:**
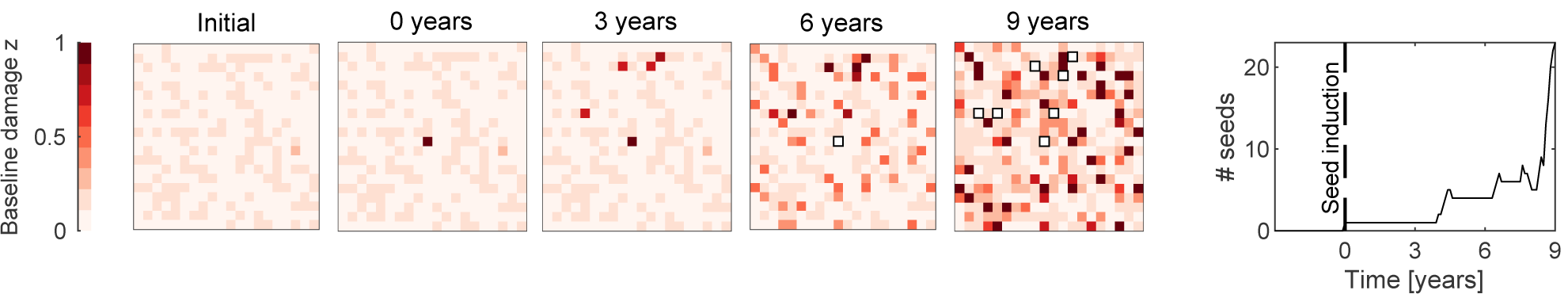
Exponential spread of baseline damage *z* after seed induction. At time *t* = 0, a seed is induced in an otherwise healthy tissue of 20 × 20 cells with mean baseline damage *z* = 0.05 ± 0.02. The damage spreads from the seed with timescale *τ*_∞_ = 10 years (see Eq. 1). Each cell receives damage from *M* = 10 other cells. Here, cells become seeds themselves if *z* > 0.5, and they die if *z* = 1 (white squares). Right panel: Damage progression is characterized by a phase of slow subthreshold spread, followed by an exponential increase in the number of seeds. Note that the spread scales linearly with *τ*_∞_ in time, and is therefore qualitatively independent of the choice of *τ*_∞_.

One effective way to avoid this spread is the quick removal of individual seed cells. The model therefore implements two fast mechanisms to maintain long term tissue health: targeted removal of individual seed cells, and quick repair of subthreshold stress in cells (see e.g. Ref. [26, 27] and Discussion for further cellular analogies). We then ask the question how and under which circumstances such a removal process can fail. The results can provide insights into what might be happening before a disease outbreak. For this, a useful seed removal model must only contain a very limited number of assumptions and parameters. The model proposed here (see Fig. 2) is built on the following pillars: (1) First, we assume seed removal is fast, and happens within hours (timescale *τ* = 4 hours) such that *τ* ≪ *τ*_∞_. A second state variable is introduced to capture this quickly evolving acute damage *x*_*n*_ for each cell. In contrast to the baseline damage, which represents irreversible damage to a cell, acute damage is temporary. For example, it represents short-term damage to an axonal myelin sheaths. (2) Second, the removal-process itself stems from a nonlinear feedback loop of interactions within the tissue. For example, glia react to neuronal damage by activating a spectrum of states which include anti- and pro-inflammatory phenotypes [27, 28]. We therefore model the removal process as the result of a tug-of-war between damage-repairing and damage inducing tissue forces. Here, repairing forces are linear and pull the acute damage *x* toward the baseline damage *z*. Therefore, these forces act homeostatically in the short term by pushing the acute cellular state back to its baseline state. The damage-inducing forces are nonlinear and increase quadratically with acute damage, and scale with the responsiveness parameter *G*. Here, we choose a quadratic term for simplicity to capture the increasing sensitivity, but other nonlinear terms are also possible. As a consequence of this tug-of-war, once a threshold of damage is passed, damage-inducing effects take over and drive the cell to cell death at *x* = 1 (see Fig. 3). This is a bistable state: Either the cells state is stably pulled to its baseline damage, or it is driven to cell death. The transition point between these two states is determined by the parameter *G*. If *G* is too small, then damage-inducing nonlinear effects are not strong enough, and the system fails to drive a cell with high baseline damage to cell death (see Fig. 3 a). But if *G* is large enough, then nonlinear damage-inducing effects will be larger than homeostatic effects and the cell is driven to cell death. Taken together, such a removal mechanism with adequate choice of *G* quickly removes seeds from the tissue, and stops the slow spread of damage in its tracks. Of course, the tissue is an assembly of cells which do not act independently from each other, and this needs to be considered in the model.

**Figure 2:**
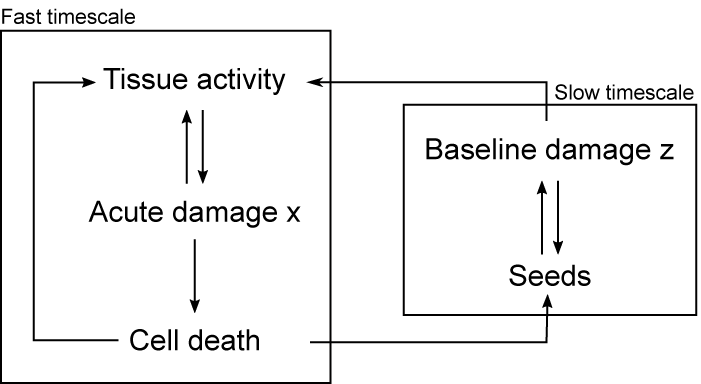
Model setup: The model consists of slow and fast dynamics. Seed-driven damage spread is modeled by an increase in baseline damage at slow timescales (see Eq. 1). Repair-or-dispose decisions are modeled at fast timescales (see Eq. 2) as an interplay between acute damage and baseline damage. Once acute damage crosses a threshold, cell death is induced, which sets off a protective effect on neighboring cells.

**Figure 3:**
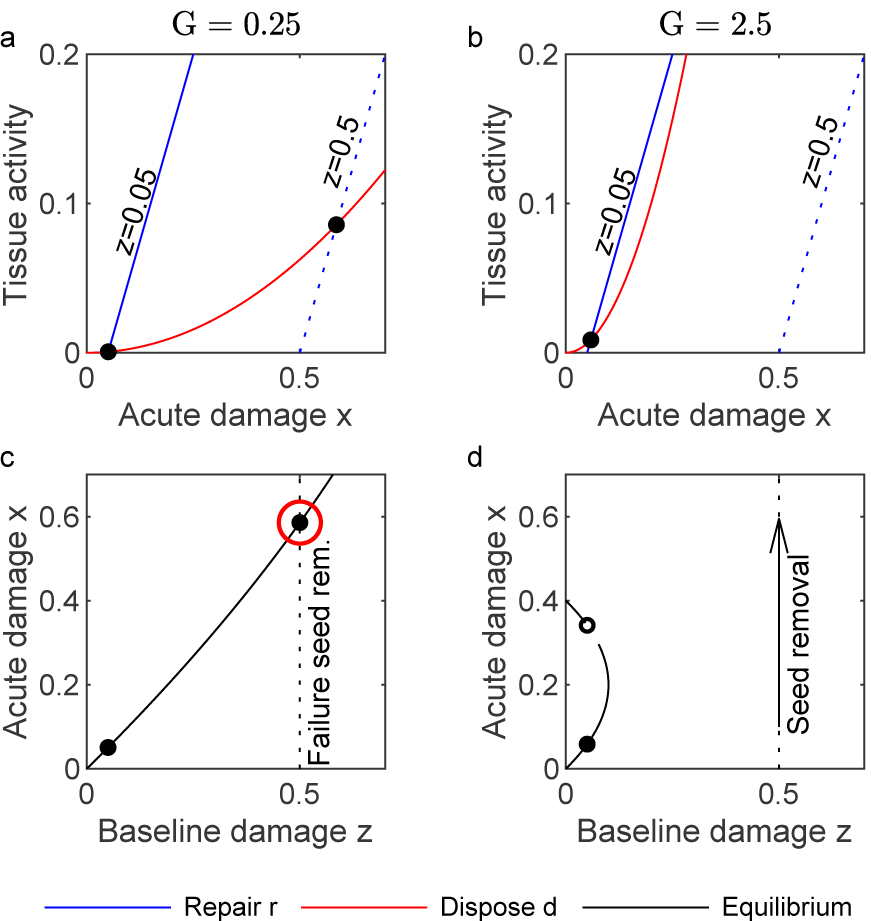
Repair-or-dispose decision. **a**,**b:** Effects inducing repair (*r* = *x* − *z*, blue lines), and effects inducing disposal (*d* = *Gx*^2^, red lines) for a system of a single cell, for scenarios with weak and strong responsiveness *G* and low and high baseline damage *z*. Black bullets show equilibrium states. **c**,**d**: Equilibrium states as a function of baseline damage *z*. The system with low responsiveness *G* maintains the seed (*z* = 0.5) at elevated levels of acute damage. The system with high responsiveness *G* does not exhibit an equilibrium state for elevated baseline damage levels, and drives these cells to cell death (‘seed removal’) (see also Sec. S1).

Cell-signaling mechanisms and cell migration [29, 30] mean that neighboring cells are impacted by the seed removal mechanism and by cell death [31]. In the ideal case, cell signaling implements a repair-or-dispose decision which is perfectly tailored for each individual cell. If this is not the case, then the tissue response may be ‘out of focus’. Here, the degree to which the damage-inducing effects are out of focus are captured by the focal range *c. c* → 0 represents perfect focus, i.e. no spatial coupling, while larger values implement a less precise damage-inducing tissue response. The effect from neighboring cells does not need to be damage-inducing. If a cell dies, then it typically has a protective effect on its neighbors (see Discussion for details). Here, we assume that this effect decays exponentially with time after cell death with constant *τ*_death_ = 24 hours, and that it scales with *D*. The full seed removal mechanism is formalized by (see Fig. 2):

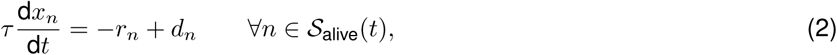

where 𝒮 _alive_(*t*) is the set of alive cells at time *t*: 𝒮 _alive_(*t*) = {*n* : *x*_*n*_(*t*′) < 1, ∀ *t*′ ≤ *t*}. The first term *r*_*n*_ = *x*_*n*_ − *z*_*n*_ represents (short-term) homeostatic forces that peg the acute to the baseline state. The second term samples damage-inducing and post-cell death effects from the neighborhood of the cell,

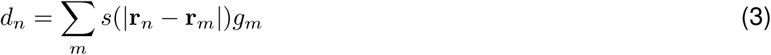

where 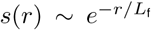 decays exponentially in space and **r**_*n*_ is the position of cell *n* in space. To allow for visual inspection and to keep the model simple, we consider *n* cells as discrete units organized on a periodic equidistant lattice with grid spacing unity. Computations are performed on one- and two-dimensional lattices. We use the focal range 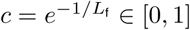 as a measure for the spatial coupling. *g*_*n*_ are effects induced by neighboring cells:

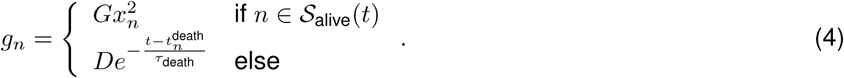

For details on the model see Methods. Fig. 4 shows a typical seed removal process. After seed-induction, damage-inducing tissue activity causes acute damage to both the seed, and its surrounding tissue. This response of neighboring cells slows down the response similar to a form of inertia. Once a threshold is passed, acute damage accumulation accelerates and the seed is driven to cell death. At this point, protective mechanisms are activated, and the surrounding tissue is ‘cooled down’.

**Figure 4:**
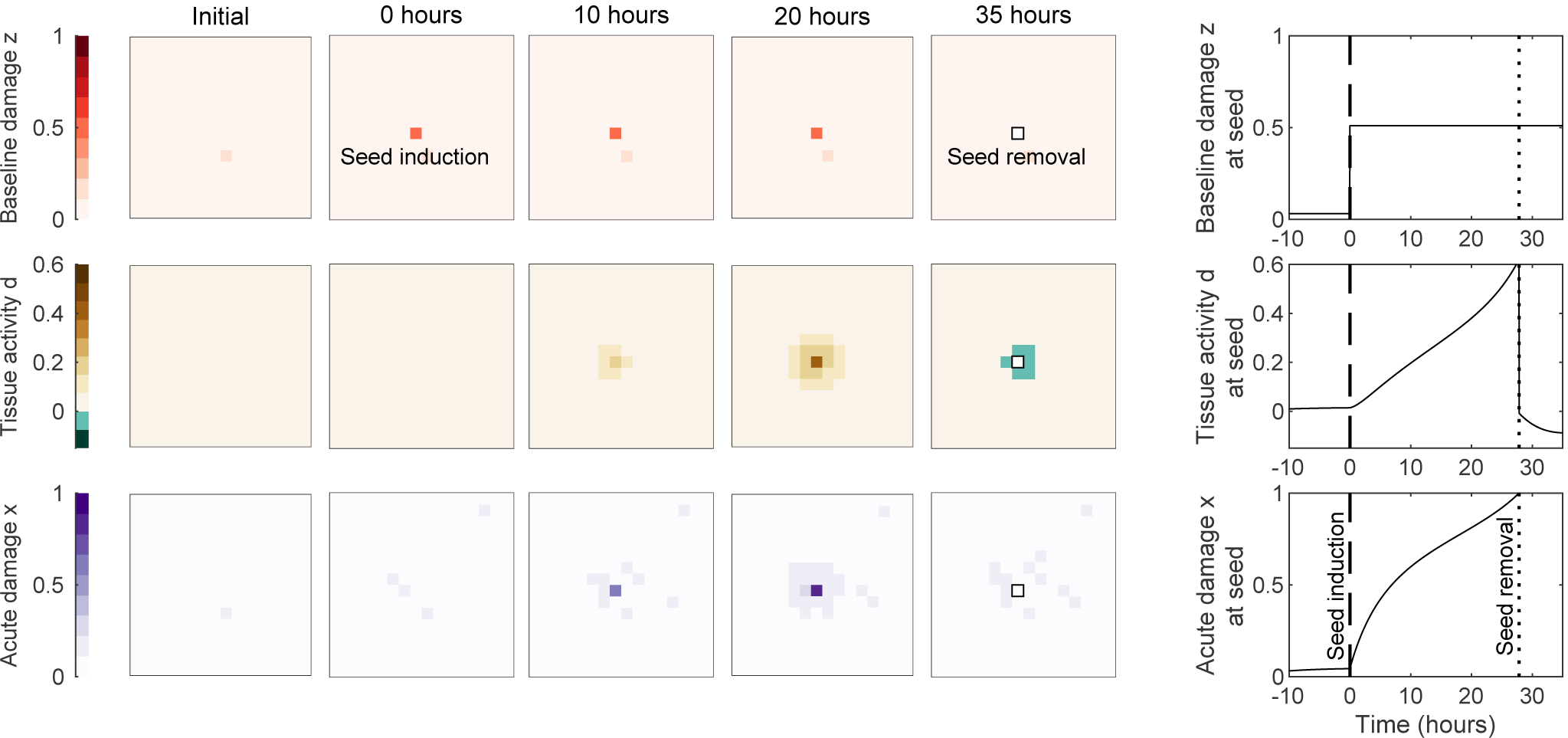
Seed induction and disposal in a healthy tissue (baseline damage *z* = 0.05 ± 0.02). After seed induction at *t* = 0 hours at the center, the tissue increases disposal-inducing effects, due to its high responsiveness *G* = 3. Elevated levels of focal range *c* = 0.4 mean that surrounding tissue is affected, too. The seed is subsequently driven to cell death (*x* = 1 at *t* = 28 hours), after which protective effects for the surrounding tissue are activated (*D* = −1, see Eq. 4), and the remaining tissue returns to its homeostatic set point.

### Tissue states

Seed removal prevents long-term damage propagation as in Fig. 1 and is therefore crucial to maintaining long-term tissue health. A systematic study of seed removal in a cell assembly revealed that there exist four states: healthy tissue, challenged tissue, primed tissue prone to acute lesion spread, and chronically inflamed tissue (see Table 1). Each state is characterized by a signature of the four model parameters: baseline damage *z*, tissue responsiveness *G*, focal range *c* and reaction to cell death *D*. It is the combination of all four parameters that determines whether seed removal is successful or not. Consider first a healthy tissue, with low baseline damage.

**Table 1:**
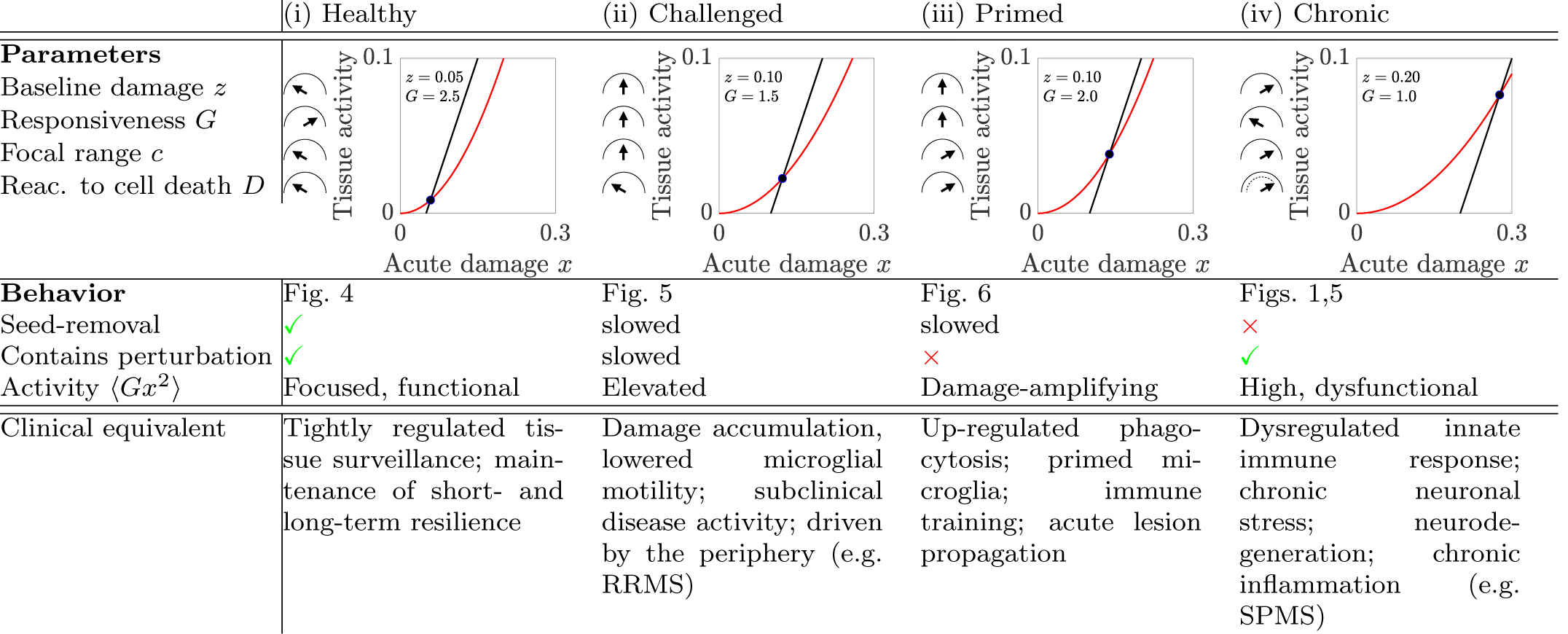
Four characteristic states of tissue resilience. (i) In the healthy state, *z, c*, and *D* are low, and high responsiveness *G* enables seed removal, therefore maintaining long-term resilience; (ii) under challenging conditions, i.e. increased baseline damage *z* and focal range *c*, the system becomes less stable, and seed removal gradually becomes less functional with decreasing responsiveness *G*; (iii) if responsiveness *G* does not decrease, the system moves closer to a critical transition, and an acute attack can induce a quickly spreading lesion; (iv) in a highly damaged system, responsiveness is down-regulated to ensure short-term stability. Nevertheless, high damage induces high levels of dysfunctional tissue activity. Insets show levels of damage-inducing (red lines, *Gx*^2^) and damage-repairing (black lines, *x* − *z*) tissue activity as a function of acute damage *x*. The steady state is shown by a black dot at the intersection of both lines.

In healthy tissue, the self-cleaning mechanism is intact and seeds are swiftly detected and removed (see Fig. 4). Ideally, this state is characterized by low baseline damage *z*, high responsiveness *G*, a narrow focal range (low *c*), and a protective tissue reaction to cell death (low *D*). Shortand long-term resilience are maintained. In this state, the tissue detects single seeds and removes them with minimal effect on surrounding tissue. It therefore prevents the long-term spread of baseline damage. Also, its high responsiveness *G* means that seeds are removed quickly. This process can, however, also be inhibited.

An inhibited, or slowed-down seed removal process is the hallmark of the challenged state. Challenged tissue exhibits increased values of baseline damage *z* and focal range *c* and requires a down-regulation of tissue responsiveness *G* in order to maintain short-term stability. Notably, seed removal is still functional in this state. It is, however, slowed down and perturbations of the tissue are resolved at a slower pace (see Fig. 5 b). In particular, an increase of the focal range *c* has dramatic effects for seed removal. At first, removal is slower and puts more stress on the surrounding tissue (see Fig. 5). The challenged phase is crucial for disease progression. Once the reserve of this phase is exhausted and seed removal fails, slow damage propagation takes over and the tissue transitions to the chronic phase.

**Figure 5:**
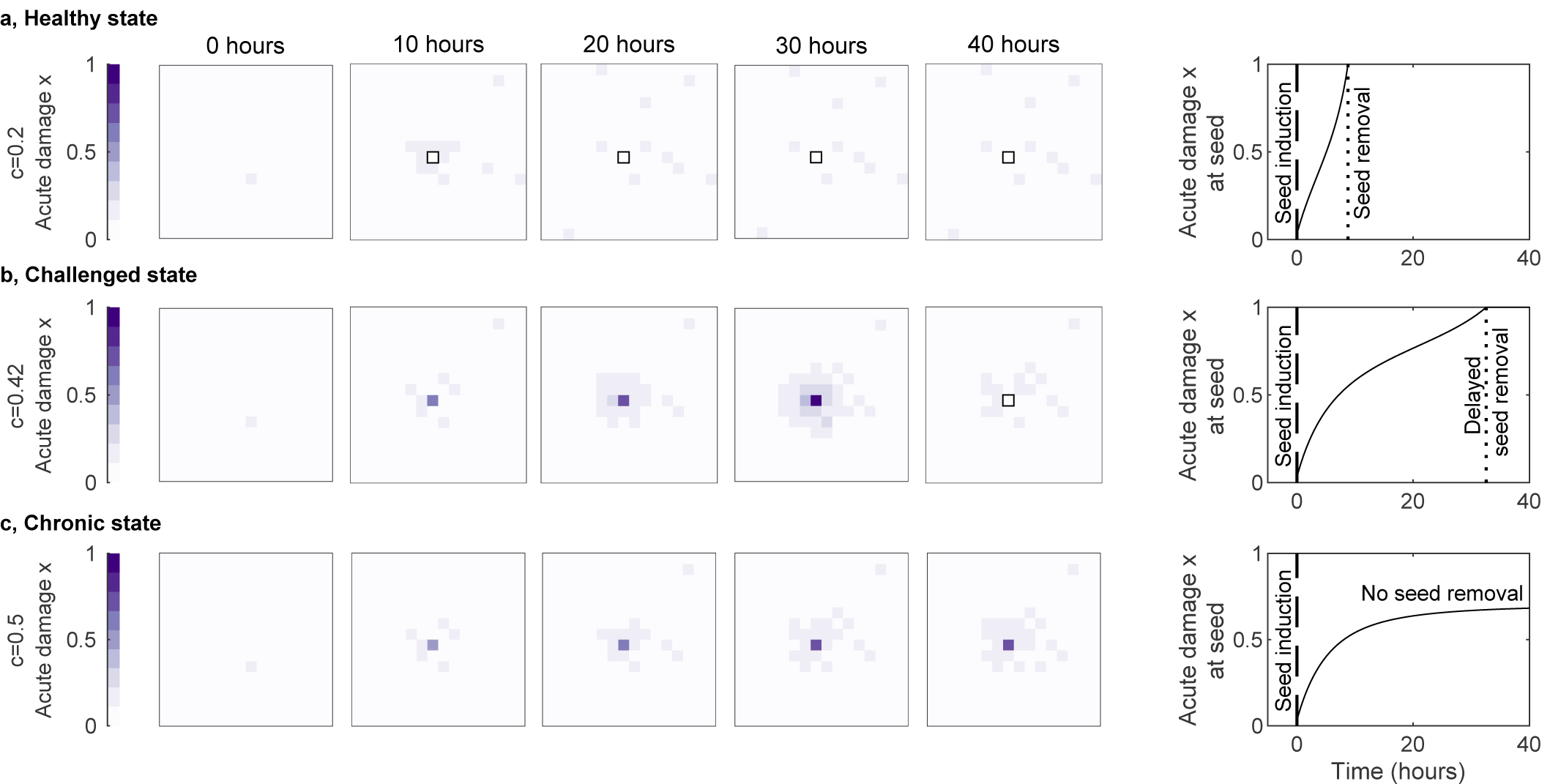
Increase of focal range *c* induces transition from healthy to challenged to chronic state. Seed induction in a healthy tissue as in Fig. 4, but with low (*c* = 0.2, **a**), increased (*c* = 0.42, **b**), and large (*c* = 0.5, **c**) focal range. Low focal range speeds up seed disposal. Increased focal range leads to delays in seed removal, and surrounding tissue is affected during seed removal. Further increase leads to failure of seed removal, and persistent stress on the surrounding tissue.

The chronic phase is characterized by a widening of focal range *c* and loss of responsiveness *G*. This means that individual cells are maintained in the tissue, even though they have very high baseline damage levels (see Fig. 5 c, Sec. S2 and Fig. S2). This failure of seed removal leads to a breakdown of long-term resilience. In this state, tissue activity *Gx*^2^ is high due to high levels of baseline damage *z* and high levels of acute damage *x*. This reflects a self-cleaning mechanism which is constantly on high alert, but is ineffective. There is a positive side-effect of this state: the down-regulated responsiveness protects the tissue against unwanted amplification of perturbations.

Such unwanted amplifications of perturbations are characteristic for the primed tissue state. In this state, the tissue appears to be healthy, but in the event of a sufficiently strong perturbation, a dysregulated self-cleaning mechanism exacerbates the effect of the perturbation, leading to a quickly spreading lesion (see Fig. 6). The primed state can be aggravated and triggered more easily if reaction to cell death does not protect surrounding cells, therefore allowing for a fast domino-like lesion propagation (see Fig. S3, S4). In the event of a large shift in conditions, the whole system may become unstable, leading to spontaneous cell death across the tissue.

**Figure 6:**
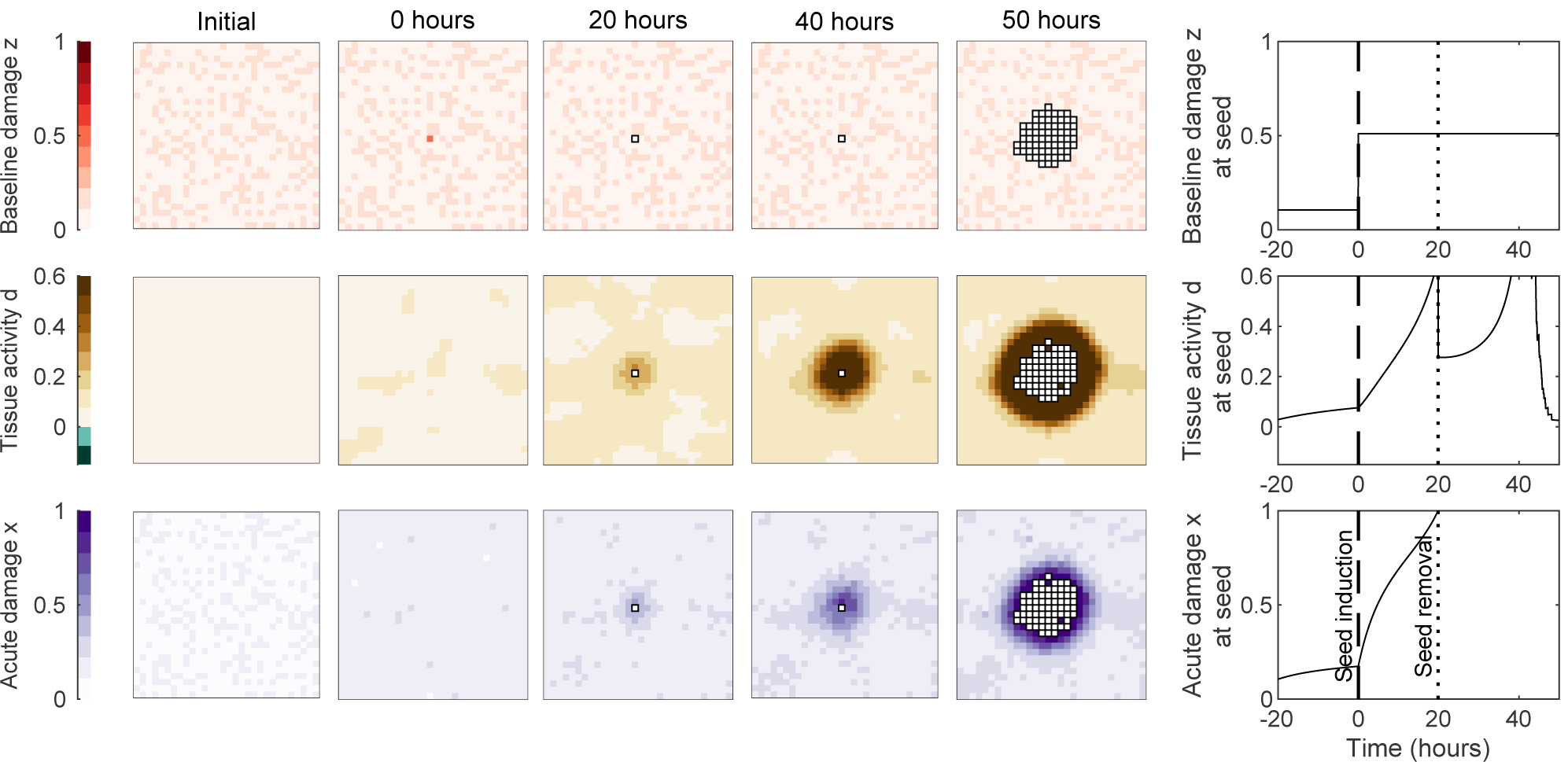
Primed tissue state leads to acute damage propagation after perturbation. Seed induction in a tissue with slightly elevated baseline damage *z* = 0.075 ± 0.035, increased focal range *c* = 0.6 and high responsiveness *G* = 3 and attenuated protective reaction to cell death *D* = 0. Disposal of the seed is slow, and affects surrounding tissue disproportionally. In contrast to Fig. 4, seed removal does not successfully inhibit disposal-inducing effects. Instead, a domino-like spreading lesion is induced, with high levels of damage-inducing activity preceding the lesion front.

We conclude that whether seed removal is functional or not depends on the interplay between all parameters. So, which possibilities exist for the tissue to adapt to increasing baseline damage levels, e.g. in age and disease?

### Tissue adaptation

We consider two stress scenarios: In a first scenario, we homogeneously increase baseline damage *z* throughout the tissue. In a second scenario, we then consider a locally constrained increase in baseline damage (see Fig. 7). We consider two measures for the tissue reaction: the time between seed induction and seed removal (see Fig. 4), and the typical time time the tissue takes to repair a small perturbation (see Methods for details). Both measures capture opposing effects: Seed removal requires a strong local damage-inducing effect to drive the seed to cell death, but the repair time hinges on repair effects being stronger than damage-inducing effects. The tug-of-war between both defines the window of allowed parameters.

**Figure 7:**
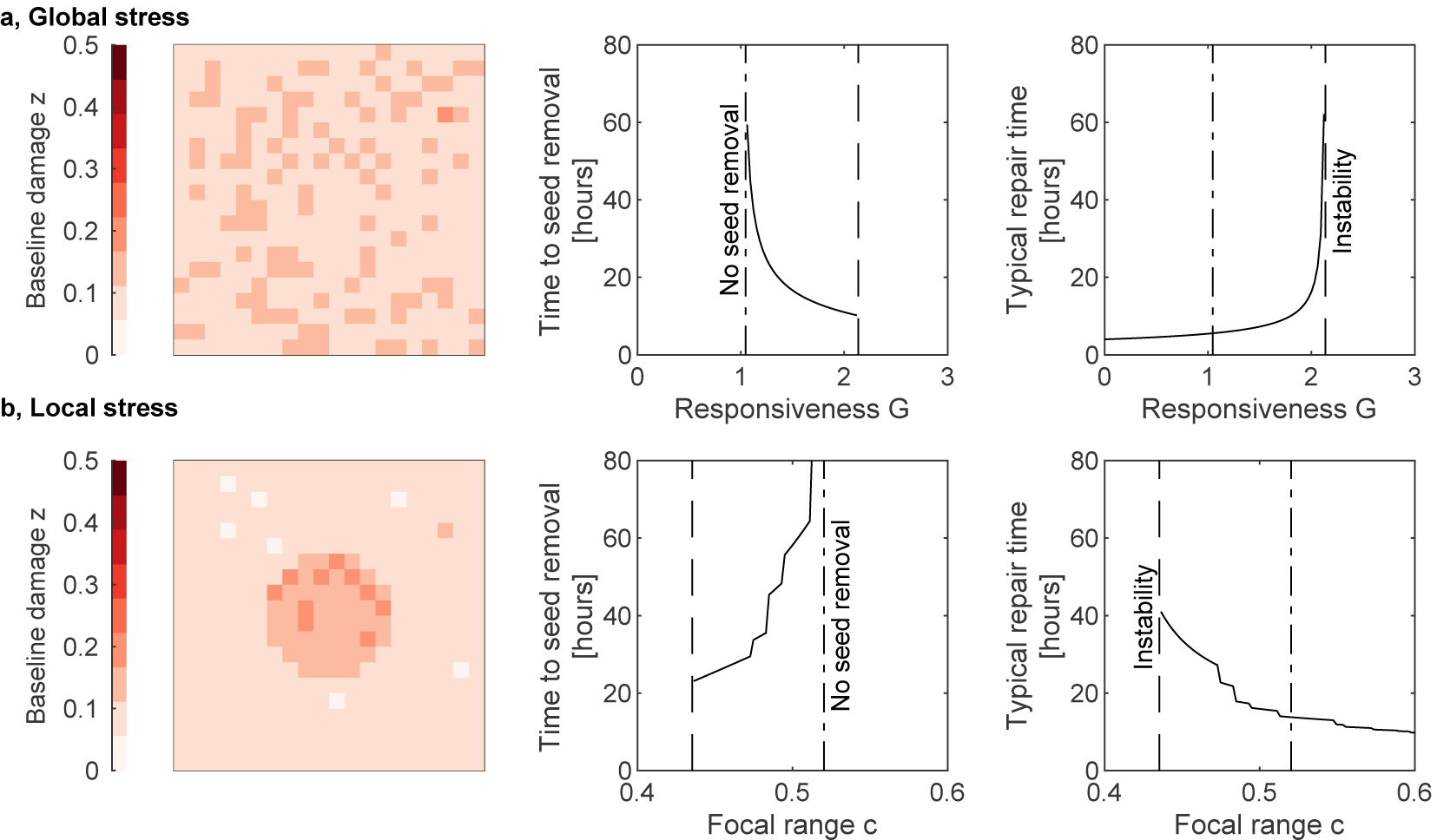
Transitions of tissue behavior. Behavior of tissue with homogeneous (**a**) and with locally constrained (**b**) baseline damage. We measure the time from seed induction to seed removal, as well as the typical timescale in which small perturbations are repaired. The latter measure is defined as the inverse of the maximal eigenvalue of the right hand side of Eq. 2. Low responsiveness *G* leads to a failure of seed removal. In contrast, high responsiveness *G* leads to an increase of typical repair time, i.e. a reduced stability at the homeostatic set point. After crossing a critical value, the set point is lost, and self-damaging effects drive cell population toward cell death. The corresponding parameter space is denoted by the term ‘instability’. For local stress scenarios, the focal range *c* can have a stabilizing effect and avoid instability. If values of *c* are too high, then seed removal breaks down. For **a**, the baseline damage is 0.1 ± 0.02, and for **b**, the local baseline damage is *z* = 0.15 ± 0.02 and otherwise *z* = 0.075 ± 0.02.

Both the homogeneous as well as the local stress scenario constrain the responsiveness *G*: A choice of *G* which is too low leads to a gradual increase of the duration of seed removal and then failure of seed removal. If *G* is too large, the repair times increase and eventually the tissue becomes unstable (see Fig. 7 a). Similarly, in the local stress scenario, if the focal range *c* becomes too large, then the tissue looses its focus and with it its ability to locally remove seeds, and transitions to the chronic state. Interestingly, strong local stressors introduce another constraint: If the focal range becomes too small, then the local stressor induces an instability. An elevated focal range therefore blurs out local subthreshold stressors and allows the tissue to remain functional (see Fig. 7 b). How do these limitations play out as stress levels increase?

We studied the two aforementioned scenarios of homogeneous and local stressors, under increasing baseline damage (see Fig. 8). The effect is dramatic – not only does an increase in baseline damage *z* shift the optimal setpoint of the tissue, therefore requiring an adjustment of responsiveness and focal range. Also, the range of allowed parameters decreases, until it eventually becomes impossible to satisfy both the seed removal and the tissue stability conditions. An increase in baseline damage also means that even with an optimal adjustment of parameters, seed removal will still slow down, and repair times will increase. Finally, a particularity of a local stressor is that the adjustment of the focal range can maintain stability. This adjustment comes with side effects such as increased reaction to subthreshold stress and possible damage propagation (see Fig. 5, 6).

**Figure 8:**
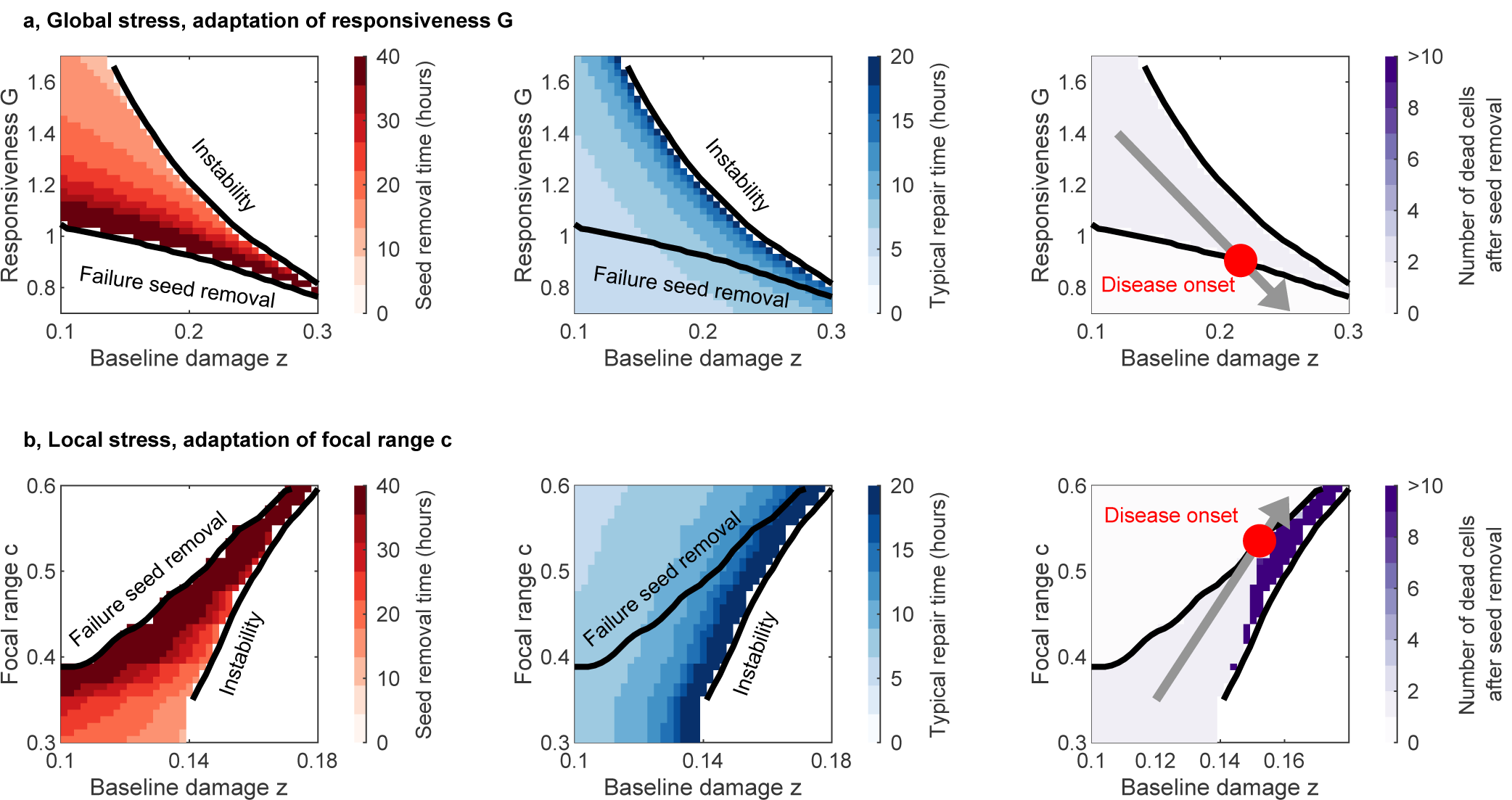
Tissue adaptation to increasing baseline damage *z*. Time between seed induction and seed removal (left column), typical repair time (center column) and number of dead cells after seed removal (right column). Transitions to instability and to failure of seed removal are defined by points at which repair time and seed removal times diverge, respectively. **a**: Global stress scenario, with adaptation of responsiveness *G* for varying baseline stress levels *z*. **b**: Local stress scenario, with adaptation of focal range *c* for varying baseline stress levels *z*. As the baseline damage increases, both the responsiveness *G* (left panel) as well as the focal range *c* (right panel), become increasingly constrained to avoid the loss of a stable set point (‘instability’) and failure of seed removal. The plots on the right depict a possible course of tissue adaptation as baseline damage *z* increases. The point at which the seed removal fails marks the onset of slow degeneration (‘disease onset’). Distribution of baseline damage as in Fig. 7. See Figs. S5 and S6 for a simulation with gradual adaptation of *G* and *c*.

In conclusion, reconciling seed removal and stability of the tissue increasingly restricts the parameter space as the baseline damage *z* increases. Maintenance of shortand long-term resilience therefore become conflicting goals for elevated baseline damage *z* levels.

## Discussion

We presented a model that describes repair and dispose decisions as a function of the tissue state. How do the model assumptions and the model behavior compare with biological observations? Can the model indeed map some aspects of the progression of complex neurological diseases? We first discuss the experimental findings and the cellular mechanisms that motivated the fast repair and dispose model. We then discuss how the model captures slow disease-specific processes. In particular, we present a re-interpretation of disease stages for Alzheimer’s and multiple sclerosis, based on the model behavior.

### The role of cellular mechanisms in seed removal

#### Repair-or-dispose

The model assumes the repair of cells with short-term-damage and disposal of highly damaged cells, as a tug-of-war between two opposing forces in the tissue. We suggest that glia-neuron crosstalk implements such a bistable effect. Glia are the helper cells of the brain, and they maintain homeostasis, and drive and resolve inflammation in the brain. The two most common glial cell types, microglia [32, 27, 33, 34, 28] and astrocytes [35, 36] exhibit differential and multidimensional activity patterns, which depend – among other factors – on the neuronal state. For instance, microglia provide neurotrophic support [37] and surveil the tissue [38]. Microglial activity is suppressed by so-called neuronal ‘Off’-signals, which regulate sensing and housekeeping functions [27, 33, 34]. These signals that induce repair are mediated by receptor-ligand interactions such as CX3CL1 and CD200 signaling [39, 37].

Glial activation is, however, a double-edged sword and can, under certain conditions, induce neuronal damage [40]. For instance, neurons may directly recruit microglia by releasing ATP, facilitating phagocytosis [41] or by releasing glutamate, which triggers microglial TNF*α* release [34]. Astrocytes respond to infections, trauma or inflammation by reactive astrogliosis, characterized by an increased production of pro-inflammatory cytokines such as IL-1*β* and TNF-*α*, and reactive oxygen species (ROS) [42]. Overactivated microglia can induce the transition of nearby astrocytes to a highly neurotoxic phenotype, causing widespread damage to the tissue [43]. Microglia can also remove stressed-but-viable neurons as a response to the ‘eat-me-signal’ phosphatidylserine following elevated, but non-toxic levels of glutamate, oxidative stress or growth-factor withdrawal [44, 26].

As a whole, these interactions between neurons and glia are high dimensional and involve a multitude of cell types and phenotypes. The model captures the following aspect of these highly complex interactions: That there exist two opposing net effects of glia-neuron crosstalk which are regulated the neuronal damage. These effects are modeled by the bistable nature of the balance in Eq. 2. The parameter *G*, which we call tissue responsiveness describes how readily dispose-inducing effects are activated. Therefore, a sufficiently high value of *G* describes that cellular communication is intact.

#### Spatial effects

The model also assumes that glia do not act perfectly precise in space, and may therefore affect neighboring parts of the tissue. This can happen through different pathways: For instance, pro-inflammatory phenotypes in neighboring cells can be induced by cytokine diffusion. Microglia migrate to regions of neural injury [29, 30] and form clusters as a reaction to blood-derived fibrinogen [45]. Therefore, inhibition of microglial motility [46] can lead to a loss of specificity and loss of precision. Also, astrocytic gap junction communication has been shown to play a role in spatial propagation of cell injury [47], and neuroprotection in ischemia [48, 49] (see [50] for a review). Any inhibition, or dysregulation of these spatial effects is modeled by an elevation of parameter *c* in Eq. 3.

#### Glial reactions to cell death

An important component of seed removal is the down-regulation of disposal-inducing effects after seed removal, i.e. after cell death. This assumption is supported by several observations. Cell death and phagocytosis is a highly complex and heterogeneous process [51]. In this process, microglia clear neuronal debris [32], secrete anti-inflammatory cytokines and minimize damage to surrounding cells [31, 52, 53]. These protective effects are captured by negative values of the parameter *D* in Eq. 4. It is not fully clear how long this process takes. However, ‘phagocytic ingestion’ was reported to take from 15 up to 160 minutes [54, 55]. After engulfment, the apoptotic cargo must then be processed [56]. In our simulations, we therefore assume that the process from detection of the apoptotic cell, engulfment, to processing of the cargo is defined by the timescale of one day.

The reaction to cell death can be disturbed in several ways. For example, it was shown that an epigenetically mediated dysregulation of the tightly controlled clearance function can lead to damage in healthy neurons [57]. Also following necrosis [58], in age [59], and in neurodegenerative diseases [60] phagocytosis is reduced or can induce pro-inflammatory reactions via ATP released from dying neurons [61]. In particular, necrosis can have detrimental effects on neighboring cells. It is characterized by a loss of cell membrane integrity and the uncontrolled release of cell products into the extracellular matrix following acute cellular injury caused by external toxic factors such as TNF-*α*. A programmed form of necrosis is necroptosis, also dubbed ‘cellular suicide’. In multiple sclerosis, oligodendrocytes have been found to undergo necroptosis [62]. In AD, neuronal loss due to necroptosis has recently been reported [63]. The net detrimental effects of cell death to the surrounding tissue is captured by higher levels of *D*. The model computations predict acute lesion spread as a possible outcome, see Fig. 6.

### Neurodegenerative disease progression

The model predicts the resilience of a self-cleaning tissue under stress. This situation is found in many neurodegenerative diseases: The self-cleaning properties of glia-neuron crosstalk are put under stress by amyloid accumulation, immune attacks from the periphery and cardiovascular stress. In parallel to these effects that strain glia-neuron crosstalk, the drivers of neurodegeneration are included as slow processes, that typically operate on the timescale of months to years: The accumulation of non-repairable baseline damage such as intracellular tau accumulation [64] or permanent demyelination [65] are represented by the variable *z*. The increase and spread of this damage is represented by the number of cells *M* the damage spreads to (see also Sec. S3). The spread could be via intracellular ‘protein condensation’ [66, 67], self-propagating protein assemblies [68, 69] and the vicious cycle of deleterious inflammation and neurodegeneration [70, 71]. As shown in Fig. 8, the model predicts warning signs that precede the breakdown of functional glia-neuron crosstalk and the take-over of slow neurodegenerative damage spread. In this section, we explore these analogies to the disease courses of Alzheimer’s disease and multiple sclerosis (see Fig. 9). We postulate that there are common system-level mechanisms which precede the clinical outbreak of the diseases. We note that these mechanisms exhibit similarities to processes in tumor development. For example, most clinically detected cancers must have evaded system-mediated immune responses [72], and pre-treatment tumor T-cell signatures can predict clinical responses to therapy [73]. The importance of the initial tumor microenvironment for the ‘turning point’, i.e. the development of an exponentially growing tumor is reminiscent of the dependence of the onset of neurodegeneration on glia-neuron crosstalk that we describe here for neurological diseases.

**Figure 9:**
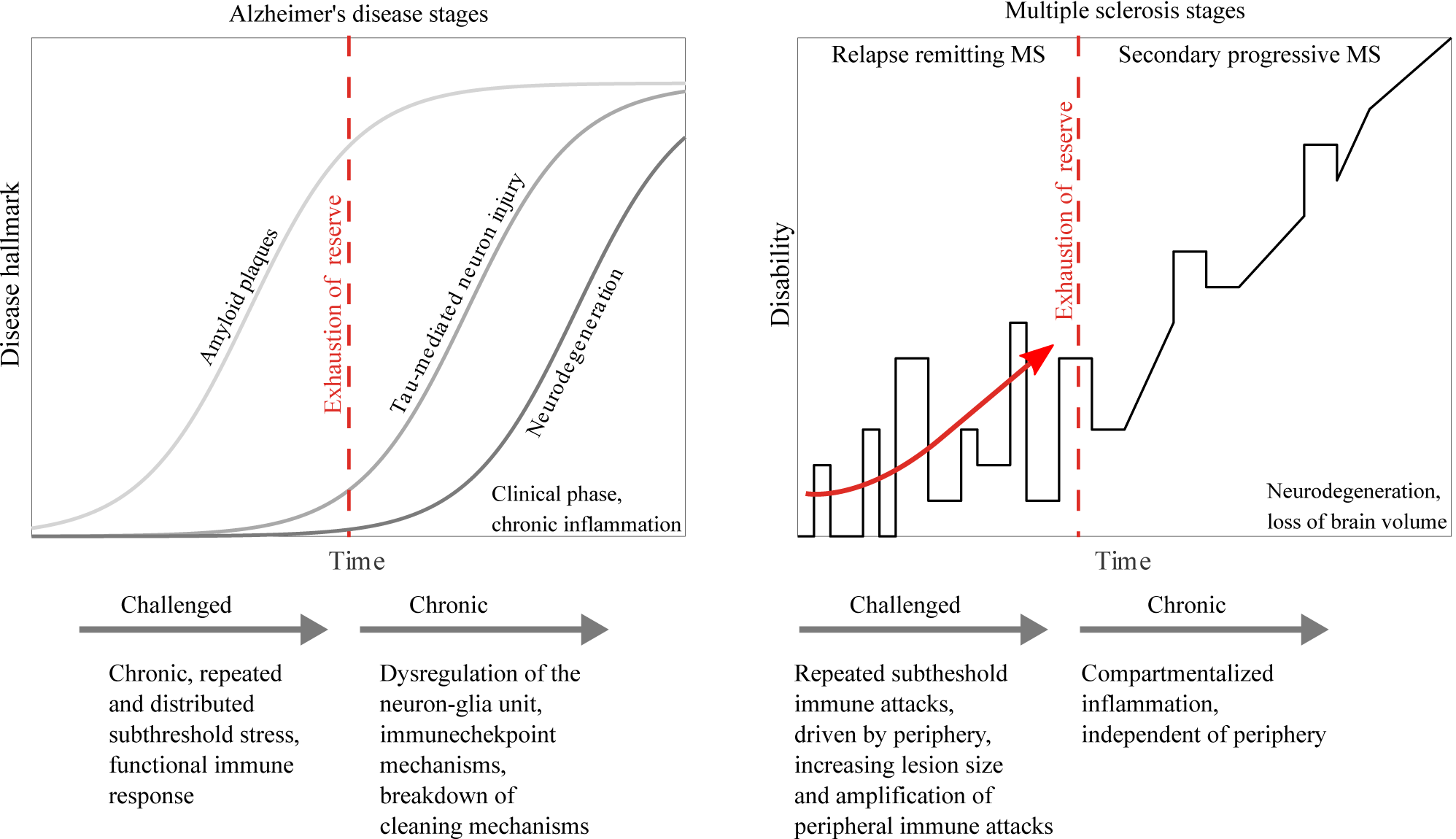
Suggested disease-model analogy. **Left, Stereotypical stages of AD disease progression:** Amyloid plaque deposition can be viewed as chronic distributed stress on the tissue, which leads to a slow adaptation. Once the reserve of this phase is exhausted, self-cleaning mechanisms break down – allowing for the spread of baseline damage, e.g. via tau-mediated neuronal injury. Only in the final phase is an increase in cell death observed. The model suggests that the breakdown of the glia-neuron unit and of functional immune responses are at the core of the transition from the challenged to the chronic and final neurodegenerative phase. **Right, Stereotypical stages of MS disease progression:** Repeated local immune attacks, driven by the periphery, are analogous to local stressors. As the limit of adaptation is approached, the effect and lesion sizes of peripheral immune attack increase (see red arrow). Breakdown of functional immune responses signals the transition from relapse-remitting MS to secondary progressive MS, i.e. the transition to the chronic phase, characterized by compartmentalization, self-driven neurodegeneration and chronic dysfunctional inflammation.

#### Alzheimer’s disease (AD)

AD is a neurodegenerative disease characterized by the extracellular accumulation of amyloid-*β* (A*β*) plaques, as well as intracellular accumulation of neurofibrillary tangles consisting of misfolded, hyperphosphorylated tau protein [74]. Next to genetic risk factors [75], also lifestyle, cardiovascular disease [76] and meningeal glymphatic system dysfunction [77] contribute to disease outbreak. It is thought that seeds recruit their soluble counterparts in living cells, therefore establishing a self-driven process. There is also evidence that neurodegeneration and glial reaction to it form a positive feedback [78, 70].

In vitro studies for AD show that A*β* induces inflammatory microglial responses [79, 80] and primes microglia for a secondary stimulus [81]. These activated glial cells appear before the first structural changes in the tissue [82]. In the early stages of aggregation the immune response amplifies the glial clearance abilities [83]. However, prolonged exposure to excessive concentrations of proteins causes chronic inflammation. Overactivated microglia and reactive astrocytes then express neurotoxic factors [84, 85, 86] and impaired A*β* clearance [87] and lose their cleaning abilities [88]. This winds up the protein aggregation [89] and compromises the inflammation resilience in a vicious feedback loop. In AD, the feedback has recently been linked to microglia- and inflammation-driven cross-seeding for A*β* pathology [68]. These processes extend the ‘amyloid cascade hypothesis’, where A*β* is seen as the main driving force of the disease. In particular, it has been proposed that AD transitions from a reversible disease course to an irreversible chronic autonomous cellular response. In this phase, disease progression no longer depends on aggregated proteins which trigger the initial response, but on glial chronic inflammation [90, 89]. We therefore include A*β* pathology as a baseline damage in our model, from which a dysregulated tissue response and failed removal of seeds ensues.

Several aspects of the role of glia-neuron crosstalk for AD progression are captured by the model proposed in this work. First, the affinity of a tissue to induce intracellular misfolded tau protein is represented by rate with which seeds are induced. Low subthreshold stressors such as lifestyle, glymphatic system dysfunction and amyloid plaque accumulation correspond to weak distributed frequent stressors on the tissue, which elevate baseline damage levels ⟨*z*⟩ (see Fig. 7, 8). Additionally, there is evidence that misfolded proteins, in particular tau and *α*-synunclein, can propagate from neuron to neuron [91, 92, 93, 24, 25] in a ‘prion-like’ manner [69]. This self-propagating seeding process may spread via synaptic connections [91, 94, 24] or other mechanisms [92], corresponding to spread of baseline damage. Another process through which the disease could spread is network dysfunction, in particular amyloid-dependent hyperexcitability of neurons with baseline activity, setting off a vicious circle of increasing neuronal activity [95, 96].

The model computations in Fig. 8 show how the slow accumulation of subthreshold baseline damage requires an adaptation of responsiveness *G*, and eventually leads to the breakdown of self-cleaning, i.e. the ability to react to intracellular tau. After breakdown of self-cleaning, the system transitions from the challenged to the chronic phase, with first an accumulation of neuronal damage and an increase in seeds (increase in tau and further increase of amyloid), and finally neuronal loss – which is in line with the typical time course of AD (see Fig. 9).

The implication of adaptive self-cleaning processes in the breakdown of resilience is in line with recent genetic studies. In particular, they implicate mutations in genes expressed by immune-active cells in the development of late-onset AD – one major risk factor being a mutation of TREM2 (triggering receptor expressed on myeloid cells 2) [39]. TREM2 encodes an innate immune receptor expressed by microglia and is part of an immune checkpoint which controls the activation of A*β*-clearing disease-associated microglia. In particular, functional TREM2 leads to an up-regulated microglial phagocytosis of A*β* plaques early in the disease by activated microglia, but it also increases levels of ApoE (plaque-associated apolipoprotein E) in A*β* plaques, which promotes misfolded protein aggregation [97, 74] – analogous to the ambiguous role of tissue responsiveness *G* in the model. In contrast, TREM2 loss-of-function mice exhibit microglia which appear to be locked in a homeostatic state [97, 98, 39]. This promotes plaque growth early but not late in the disease. This means that intriguingly, the function of TREM2 signaling is analogous to the responsiveness *G* in the model – it participates in active seed removal in healthy tissue, but is counterproductive in the chronic phase.

This hints at a possible transition from a challenged but functional phase to a chronic phase, corresponding to the so-called ‘cellular phase’, in which clearance mechanisms break down [89]. Once the chronic phase is entered, several slow neurodegenerative processes drive disease progression. For instance, dying or damaged neurons release so-called danger-associated molecular patters (DAMPs). These can initiate microglial NLRP3 inflammasome activation, enhance amyloid seeding and promote tau hyperphosphorylation [39, 37]. Experiments also indicate that A*β* plaques disseminate throughout the brain, even though the underlying mechanisms are not yet fully understood [69].

Importantly, the model suggests that the transition to the chronic phase is a self-protective measure: Consider a situation in which stress on neurons with a selective vulnerability to protein aggregation [67] becomes excessive, or if average damage levels are too high, eg due to cardiovascular disease. In this case of high baseline damage *z*, an active removal of seeds can induce collateral damage and becomes counter-productive (see Fig. 6). The model suggests that down-regulation of glial responsiveness *G*, which compromises long-term resilience at the benefit of maintaining short-term resilience, is then the lesser of two evils. Similarly, the intracellular accumulation of misfolded proteins in AD (and also in Parkinson’s disease), influences glial reactions to cell-death *D*. Glial phagocytic function may become up-regulated and negatively affect neighboring neurons [57]; and necrosis of long-term stressed neurons can lead to further inflammation and damage propagation [71]. This can lead to fast domino-like passing on of neurodegeneration to neighboring tissue, characteristic for the primed state (see Fig. 6).

In summary, the model computations suggest that amyloid plaque accumulation, together with other factors, gradually increase baseline damage. This leads to a challenged tissue state, in which in order to maintain short-term stability, the tissue continuously lowers its responsiveness. Once the reserve of this state is exhausted, self-cleaning mechanisms of glia-neuron crosstalk break down, allowing for a spread of tau-mediated neuron-injury (baseline damage). Only in the final stage, neuron death is observed.

#### Multiple sclerosis (MS)

A hallmark feature of MS is that immune cells penetrate the brain, demyelinate neurons [99] and induce excitotoxic insults [100, 101]. MS often progresses in two sequential disease phases [65]. In the first phase, also referred to as relapse-remitting MS (RRMS), neurodegeneration is driven by immune cells entering the CNS from the periphery through the blood-brain-barrier and attacking myelin sheaths and axons. In the second phase, also referred to as secondary progressive phase (SPMS), neurodegeneration is compartmentalized and chronic. The transition point between these two phases is highly patient-specific and may either happen within a few months, decades or, as in some patients, not occur at all. The reasons for the transition and its high variability are not fully understood [102]. So, are there slow self-propagating mechanisms in MS?

We identify several self-propagating processes that can induce a vicious neurodegenerative cycle, and which correspond to the slow propagation of baseline damage in the model. (1) First, in acute MS lesions, the transporter protein levels GLT-1 and GLAST are dysregulated and therefore impaired in their ability to take up glutamate [103, 104]. This is also seen in amyotrophic lateral sclerosis (ALS) [105]. This reduction in the astrocytic glutamate uptake [106] can exacerbate the toxic increases of intracellular calcium [107] following T-cell attacks in MS. This means that the tissue becomes increasingly vulnerable to excitotoxicity and external stress. (2) Second, in age and in disease, glial processes can reduce remyelination of neuronal axons. This may be a consequence of impaired myelin clearance from an initial injury due to reduced motility, surveillance, and phagocytic activity of activated microglia with age [108]. In MS, impaired myelin debris clearance can then lead to cholesterol crystalisation, inflammasome activation and a maladaptive immune response [109, 110]. Importantly, reduced myelin debris clearance, and the resulting myelin debris in the tissue, inhibits oligodendrocyte progenitor cell differentiation into oligodendrocytes [110], further aggravating the problem. (3) Third, in MS, regulatory lymphocytes, which act through crosstalk with microglia [111], are less effective in inducing remyelination. Also, chronic inflammation and microglial activation, and increasing neuronal oxidative stress and damage of the axon-glia unit, can lead to a self-perpetuating vicious cycle [112, 113]. Recent studies even point at the possibility that decoupled pathological protein-propagation contributes to this vicious cycle in progressive MS, although research is still at an early stage [114].

Apart from the self-perpetuating pro-cyclic effects described above, the model captures the local nature of repeated immune attacks around blood vessels as local increase in baseline damage *z* (see Figs. 7, 8). Permanent demyelination of neurons is captured by the baseline damage *z*, and reversible demyelination and abnormal intracellular calcium elevations are represented by acute damage variable *x*. The tissue reaction to damaged neurons is captured by the parameter *G*. As the disease progresses, the repeated immune attacks increasingly put the responsiveness of immune competent cells to the test: Glial responsiveness to immune attacks can cause collateral damage to neighboring cells, but is needed for repair and maintenance, to avoid neurodegenerative processes. The model suggests that in order to avoid a breakdown of short-term stability and collateral damage by acute lesion spread, the focal range *c* is increased. The side-effect is a less stable tissue, in which the ‘footprint’ of immune attacks increases. The increased susceptibility is reminiscent of the increasing intensity of RRMS lesions before the transition to SPMS. In the extreme case, the acute lesion spread described in Fig. 6 is similar to the development of quickly expanding lesions in fulminant MS [115]. Eventually, the adaptation to high baseline levels leads to a breakdown of seed removal – this means the removal of highly demyelinated neurons ceases, which allows for the development of slow neurodegenerative processes. Slow compartmentalized neurodegenerative processes take over, leading to a transition to the chronic phase (see also Table 1). The model therefore suggests that the point at which focal range *c* is increased and glial responsiveness *G* is down-regulated so much that seed removal breaks down characterizes the transition from RRMS to SPMS. Postponing the transition from RRMS to SPMS therefore hinges on the careful management of immune attacks and subsequent tissue adaptation.

## Conclusion

The key result of our model computation is that maintenance of seed removal and short term resilience become conflicting goals with increasing tissue damage. This is consistent with features of disease progression in AD and MS. Therefore, even though evidence suggests that overlap of genetic susceptibility across neurodegenerative diseases is low [116], our model hints at the possibility of common system-level mechanisms which precede the clinical outbreak of the diseases. So, how can a transition from a functional tissue to a primed (failure of short-term resilience) or chronic (failure of long-term resilience) be postponed? Possible targets are: (1) maintenance of the active elements of self-cleaning, i.e. responsiveness *G* and focal range *c*; (2) maintenance of reaction to cell death (*D*) and (3) minimization of long-term subthreshold stressors that lead to an increase of baseline damage *z*.

To maintain the self-cleaning mechanism, responsiveness *G* needs to stay high, and focal range *c* low. Possible avenues to do this are the maintenance of the various checkpoint mechanisms of the innate immune system [117], as well maintenance of microglial motility [46]. Both become dysregulated in age [118]. Warning signs which signal the exhaustion of the challenged state, and a possible failure of self-cleaning, are slowed responsiveness, delayed seed removal (see Figs. 5a,8) and decreased short-term stability (see Figs. 6, 8). The model therefore suggests that changes of the dynamic tissue reaction to focal stress predict disease onset.

This constitutes the model prediction to be tested: Disease onset coincides with the breakdown of a glia-neuron control mechanism which up-regulates tissue activity following injury, and that the breakdown is preceded by two opposing observations: (1) Small, repairable injury are resolved at a slower rate (see repair time in Fig. 8) and (2) injury which require an inflammatory response (seed removal) take longer to initiate that necessary tissue response (see ‘time to seed removal’ in Fig. 8). The disease onset coincides with breakdown of (2), i.e the ability to cause adequate tissue responses. Confirming or refuting this process hinges on the development of biomarkers for the dynamic tissue response to physiologically small and focused injury (seed inductions). If confirmed, this could be a potential diagnostic tool. Note that in contrast to early disease phases, in which high responsiveness *G* needs to be maintained, our results show that in the chronic phase, active seed removal is dysfunctional, calling for a down-regulation of responsiveness *G*, e.g. using anti-inflammatory treatments. This decreases the risk of collateral damage (see Fig. 6), and slows down feedback processes which drive neurodegeneration. Furthermore, an antiinflammatory protective tissue reaction to cell death is required (negative *D*). The exhaustion of this effect may lead to a primed state and therefore act as an accelerator and facilitator of neurological disease progression, in particular in protein-accumulating diseases. To avoid or postpone this transition, therapeutic interventions need to selectively reinforce the resolution of glial responsiveness after cell death.

In conclusion, we propose a self-consistent computational description of a self-cleaning mechanism under stress with a minimal number of parameters. Analysis and model computations suggest the existence of four stereotypical tissue states and a typical pattern for disease progression, dependent on the stressor. The ingredients of the model reflect several key properties of glia-neuron crosstalk and model predictions match current genetic and experimental findings for two of the most common neurodegenerative conditions. Mechanistically, the model suggests the following state-specific interventions: maintenance of glial fitness, responsiveness and focal range in early, pre-clinical phases; management of the ability to contain and resolve glial activation after cell death; and reduction of responsiveness in the chronic phase of the disease. Specifically, the model suggests how a failure to consolidate shortand long-term resilience might be at the heart of the onset of common neurological diseases.

## Material and Methods

If not stated otherwise, baseline neuronal damage levels *z*_*n*_ are independently drawn from a log-normal distribution, where mean and standard deviation are given in the figure captions. The spatial coupling term *s*(*r*) in Eq. 3 is normalized such that in the given domain, ∑ _*m*_ *s* (|**r**_*n*_ − **r**_*m*_|) = 1.

## Acknowledgments

This study was supported by the Max Planck Society (T.T.), German Research Foundation (DFG, SFB CRC-TR-128 to S.B.), the Hertie foundation (mylab, S.B.), the Joachim Herz foundation (Add-on fellowship, A.N.) and the German Academic Exchange Service (DAAD-scholarship, D.B.). The authors are members of the Rhine-Main Neuroscience Network (rmn2). We thank Jochen Roeper and Olga Garaschuk for their insights into possible roles of the mechanism in Parkinson’s and Alzheimer’s diseases, respectively. A.N. thanks Tatiana Engel for stimulating discussions about about the model framework and Daniela Sichwardt for proofreading the manuscript.

A.N., D.B., K.B., S.B., T.T. conceived the study; A.N., D.B. performed the research; A.N. conducted the mathematical analysis and prepared the first draft of the manuscript, A.N., S.B. and T.T. wrote the manuscript, and all authors revised the final manuscript.

## Supplemental Information

### S1 Mean field and steady state solutions

The mean field limit of Eq. 3 is defined by *x*_*n*_ = *x, g*_*n*_ = *g, z*_*n*_ = *z* constant for all *n*. In this case, if the cells are alive (*x* < 1), then *g* = *Gx*^2^, and the acute damage *x* solves 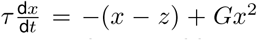, with steady states given by 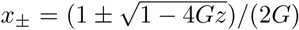 if *z* < *z*_crit_ = 1*/*(4*G*). Here, *x*_+_ and *x*_−_ are the unstable and stable solutions, depicted in Fig. 3. The critical point, beyond which no stationary solution exists, is given by *z*_crit_, see also Fig. S2.

### S2 Focal range *c*

The interplay between responsiveness *G* and focal range *c* determines the tissue reaction to cell damage. It therefore determines whether seed removal is functional or not. In Fig. S2, the population effect of four configurations of *G* and *c* is shown as an example. In the following, we study the behavior of Eq. 3 of the main manuscript in the limit of very narrow and very wide focal ranges.

If the focal range is very narrow, *c* → 0, then ∑ _*m*_ *s*(|**r**_*n*_ − **r**_*m*_|)*g*_*m*_ → *g*_*n*_, and we obtain independent ordinary differential equations for all *n*:

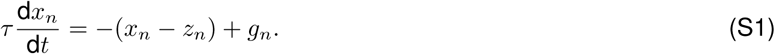

The results of the mean field section then hold individually for each cell. One side-effect of an increased focal range is that single cells can be maintained, even though their baseline damage *z*_*n*_ exceeds the critical value *z*_crit_ (see Figs. 5 and S2).

If the focal range is very wide, *c* → 1, we map the discrete values of *n* onto the spatial variable *η* = Δ · *n* with Δ = −log *c*. As *c* → 1, Δ decreases to zero, allowing us to define state variables as functions of *η*: *x*(*η*), *g*(*η*), *z*(*η*). In the one-dimensional case, *d*_*n*_ in Eq. 3 becomes

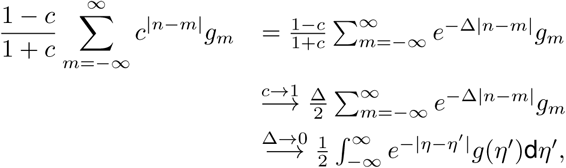

where we used that 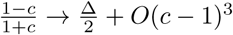 as *c* → 1. Eq. 3 then becomes an integral equation

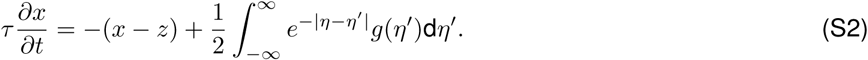

Therefore, for highly imprecise reactivities the characteristic length scale is Δ and the influence of one cell on another decays exponentially on that length scale. This can be observed in the snapshots of Fig.S4.

#### Critical transitions

High levels of baseline damage *z* can destabilize the tissue. This can happen in two ways: First, a sudden large increase of ⟨*z*⟩ can lead to spontaneous cell death across the tissue (see Fig. S3). This has different consequences, depending on the focal range *c*: Systems with narrow focal ranges allow for gradual cell loss when *z* increases, but systems with wide focal range *c* react suddenly and collectively. This population effect has been observed in many other biological and ecological systems, for instance when coral reefs are repaired by ‘mobile link organisms’ from nearby reefs [119].

As discussed in the main manuscript, in addition to this classical transition, our model also reveals a metastable ‘primed state’ for intermediate levels of *z* (see Fig. 6). Under normal conditions, this primed tissue appears to be healthy. However, in the event of strong enough perturbation, the tissue amplifies damage and may initiate a wavelike spreading lesion. In general, anti-inflammatory reactions to cell death (negative *D*) protect the surrounding tissue against these knock-on effects. If this protection, however, is attenuated, and focal range is wide, then acute attacks can cause widespread damage. Wide focal ranges (high *c*) and pro-inflammatory reaction to cell death (high *D*) increases damage. Importantly in this case, the risk of acute short-term damage amplification is more pronounced for a system with intact, high, responsiveness *G* (see Fig. S4 for details).

### S3 Seeding parameter *S*

Equation 1 describes slow damage propagation. We can add a term that represents seeding events that happen at rate *ν*. After non-dimensionalization, we obtain

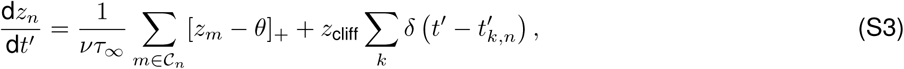

with dimensionless timescale *t*′ = *tν*, and where the seeding events 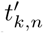 take place at a rate of one (with respect to the new timescale *t*′). The early, linear damage propagation is then described by the dimensionless parameter

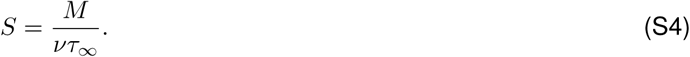

*S* represents the balance of the spreading strength *M*, given by the number of elements in 𝒞 _*n*_ and speed 1*/τ*_∞_, to the rate of independent self-induced seeding events *ν*.

**Figure S1:**
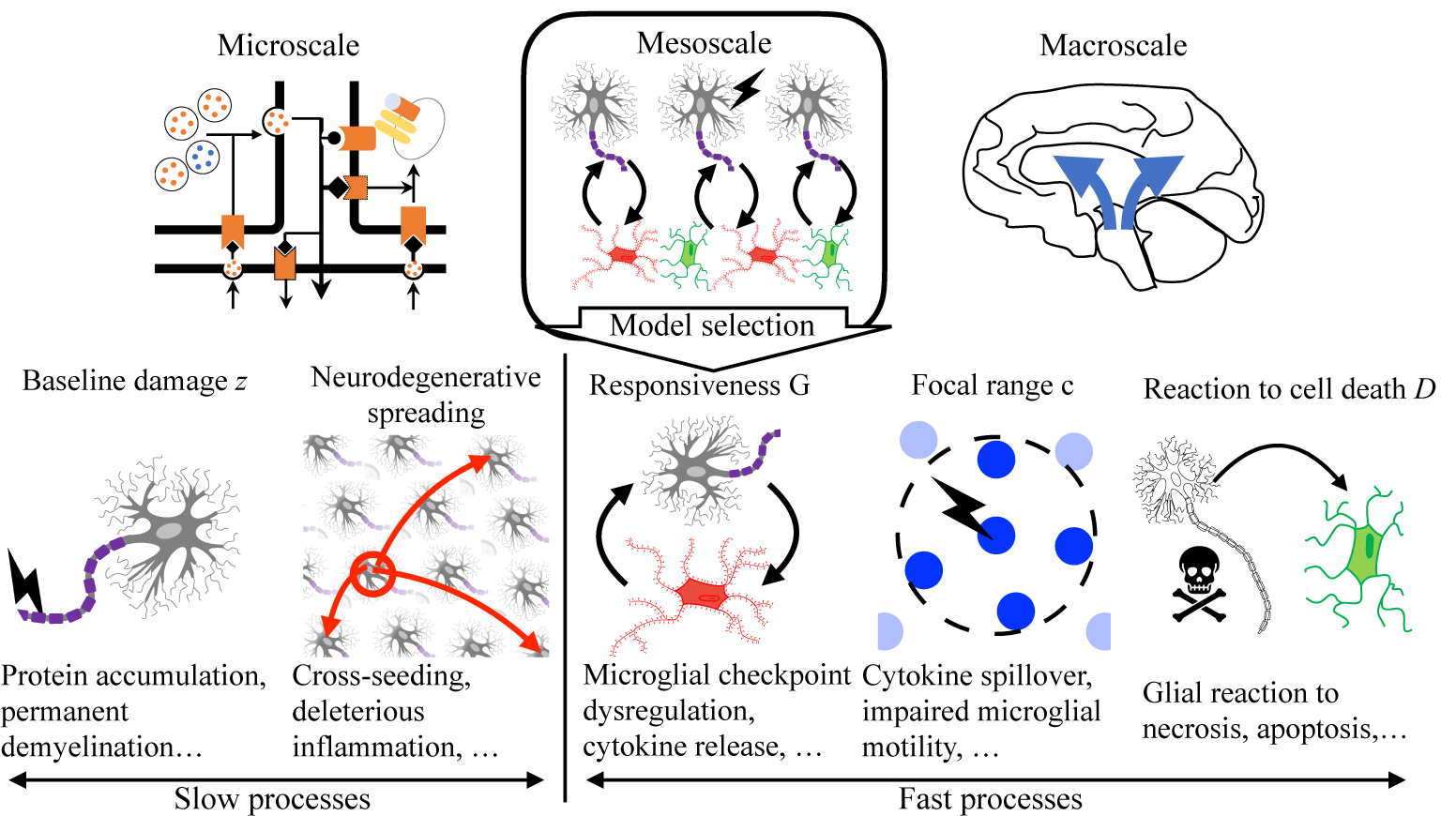
Five classes of neuron-glia crosstalk are included in a mesoscale model. We choose a mesoscopic modeling level which generalizes tissue response to a small number of observable factors. It mediates between the microscopic level of cellular and molecular processes [120] (top left) and the macroscopic level of interacting brain regions [121, 122] (top right). The model includes the following distinct properties of glia-neuron crosstalk: (i) accumulation of non-repairable baseline damage (*z*); (ii) slow neuron-to-neuron spreading of baseline damage (*S*); (iii) glial responsiveness to neuronal damage (*G*); (iv) focal range of glial reactivity (*c*); (v) tissue reaction to cell death (*D*).

**Figure S2:**
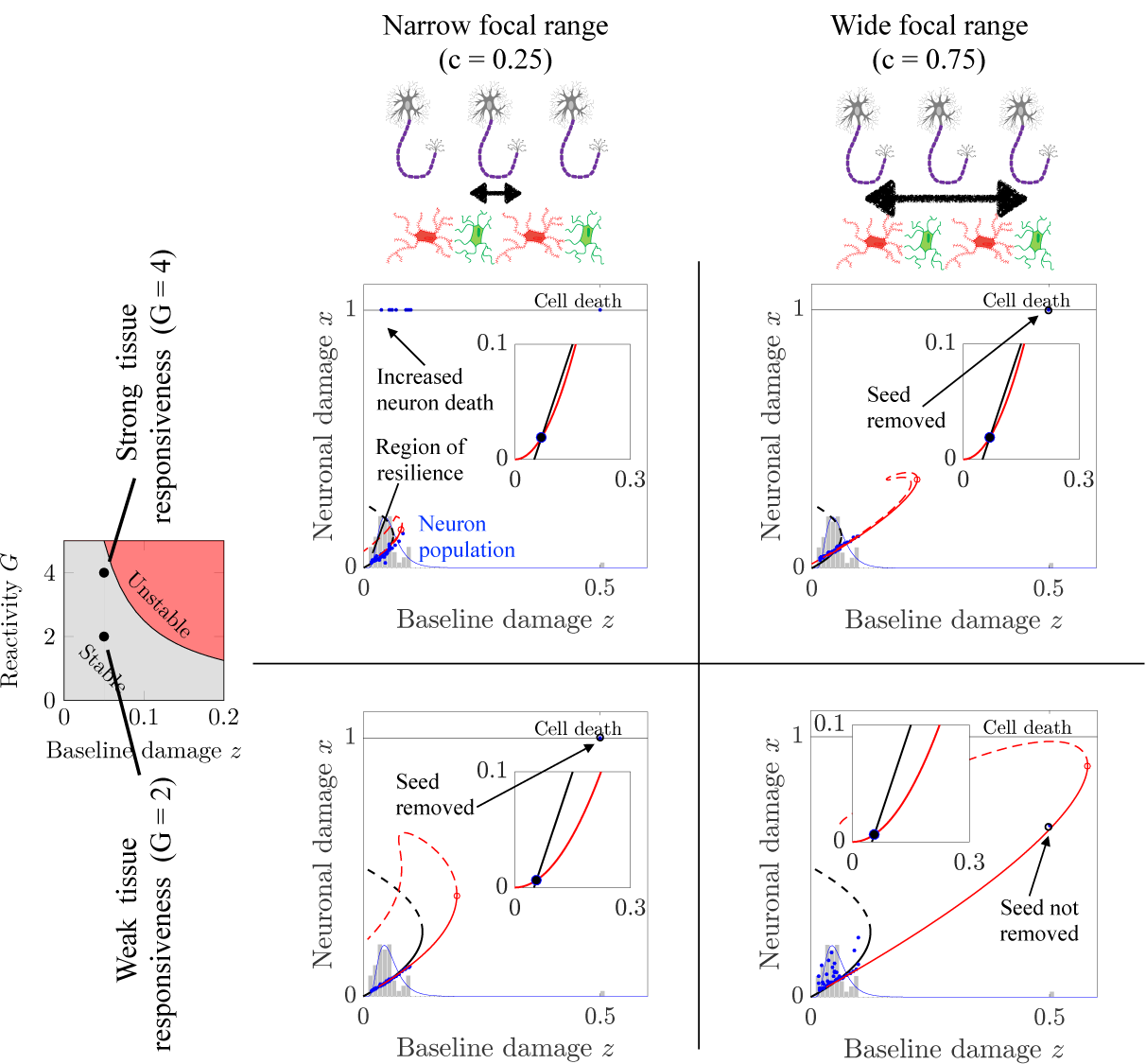
Functional tissue resilience requires tuning between focal range *c* and tissue responsiveness *G*. Black solid and dashed lines represent stable und unstable steady states in the mean field limit. Red solid and dashed lines represent stable und unstable steady states of one cell for a given (*z*_*n*_)-distribution. Blue dots represent a population of 200 cells, taken from a log-normal distribution (blue lines). Grey bars represent the histogram of their (*z*_*n*_)-distribution. Note that some cells are dead (death threshold at unity). For a narrow focal range (*c* = 0.25), cells react more independent of each other (see left column). For a wide focal range (*c* = 0.75), population effects allow for regional compensation of locally elevated stress levels (see top right), but also of rescue of seeds, if the responsiveness *G* is attenuated (see bottom right). Red and black lines of the insets show mean field damage inducing and damage reducing effects *Gx*^2^ and *x* − *z*, respectively, as a function of acute damage *x*.

**Figure S3:**
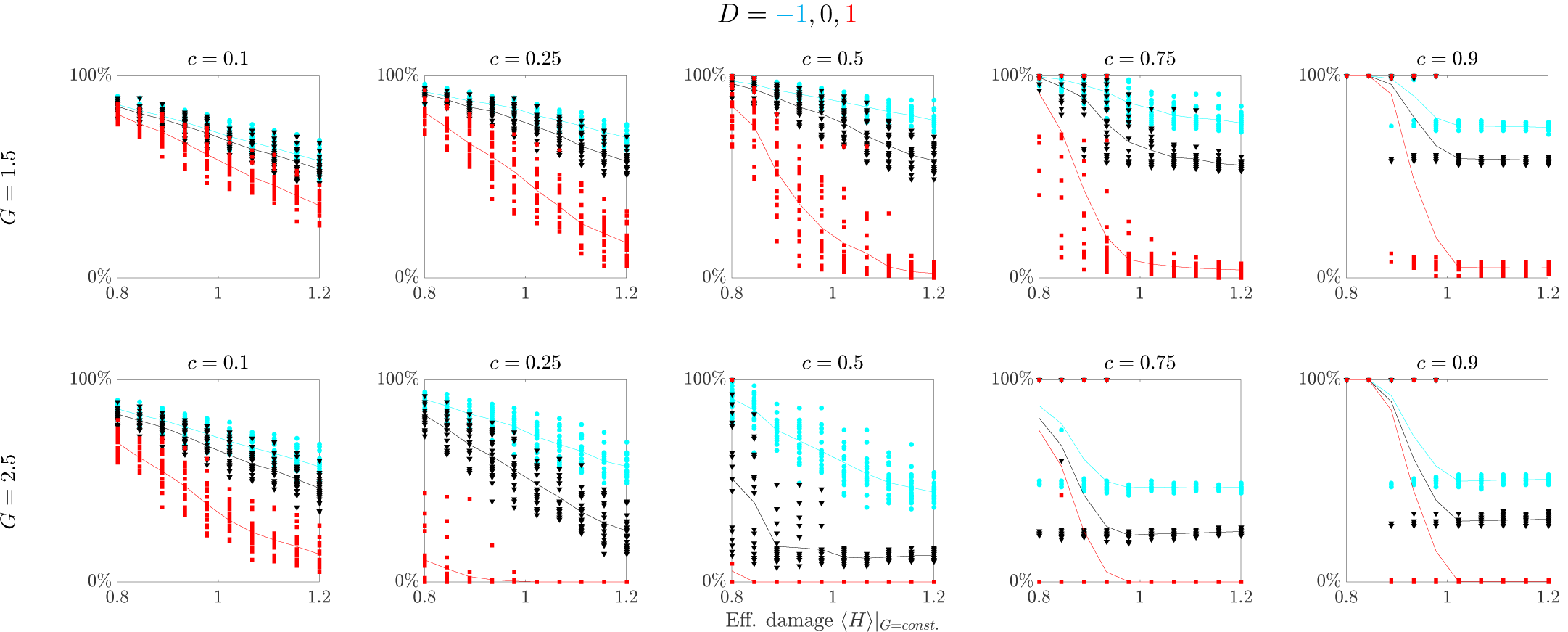
Critical transition after population increase of baseline damage. Survival rates after spontaneous cell death for different strengths of responsiveness *G* for different levels of focal range *c*, plotted as a function of the effective damage parameter ⟨*H*⟩ |_*G*=*const.*_ = ⟨4*Gz*⟩ | _*G*=*const.*_. Simulations with anti-inflammatory, neutral or proinflammatory reaction to cell death (*D* = −1, 0, 1) are represented by cyan, black and red lines, respectively. Each symbol represents one distribution of baseline damage levels (*z*_*n*_), and solid lines connect mean values. Parameters: *N* = 100 in a periodic one-dimensional domain, *t*_max_ = 10 days, 20 runs per parameter set. Systems with a wide focal range (*c* →1) exhibit step-like transitions, for which all cells survive until the critical point is reached, beyond which a steep decrease in survival rates is seen. In contrast, for narrow focal ranges (*c* → 0), the tissue reacts separately to the stress levels of individual cells, therefore allowing for a gradual decline of survival rates. Protective tissue reactions to cell death (*D* = −1) can act as a barrier for the spread of damage and stabilize cell loss after crossing the critical transition. If protective effects to cell death are attenuated (*D* = 0) or become toxic (*D* = 1), then the death of the first cell sets off a domino-like knock-on effect that can lead to the early death of large parts of the cell assembly. Tissue with intermediate focal range (e.g. *c* = 0.5) is particularly vulnerable to this effect, because local stress by a dying cell fully acts on the neighboring cell, and is not distributed across the tissue. This means that even though the system is stable locally, i.e. small perturbations to the damage levels can be compensated for, it is not stable with respect to the impact of one dying cell, which sets off a propagating lesion.

**Figure S4:**
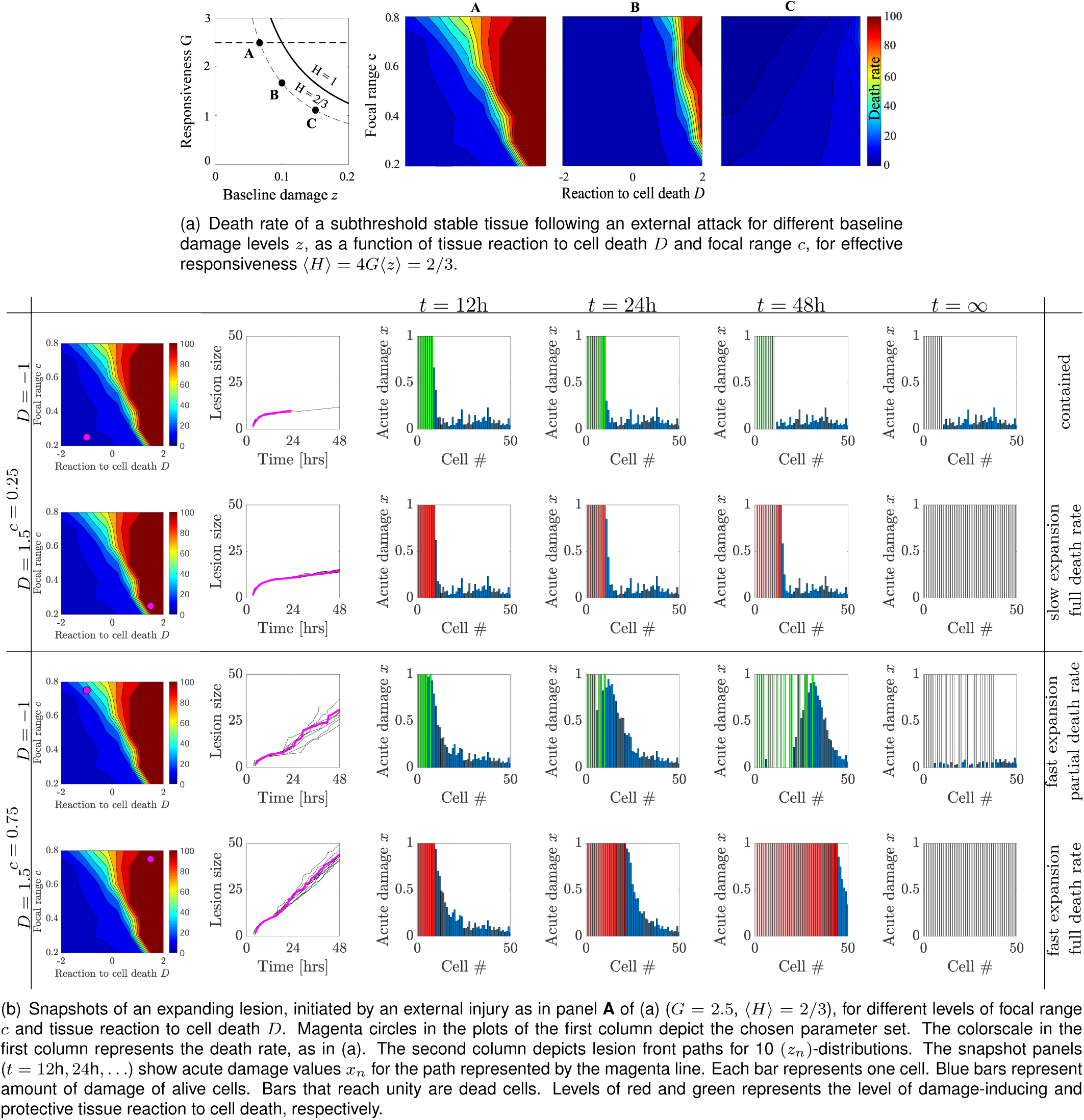
Analysis of primed tissue. Impact of an acute attack on a stable tissue with 100 cells. Here, the stressor function 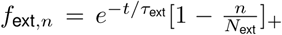, is added to the right-hand side of Eq. 3. *τ*_ext_ = 48 h and *N*_ext_ = 10. Acute attacks can initiate wave-like expanding lesions for highly reactive tissues. For a narrow focal range (*c* = 0.25), damage spreads in a domino-like manner from cell to cell. It is contained if tissue reaction to cell-death is protective (*D* = −1), but spreads slowly if tissue reaction to cell death is damage-inducing (*D* = 1). For a wide focal range (*c* = 0.75), a large enough part of the cell population needs to be sufficiently damaged to activate the toxic feedback loop. In this case, the lesion spreads quickly, leading to partial and complete cell death for protective and inflammatory tissue reactions *D*, respectively.

**Figure S5:**
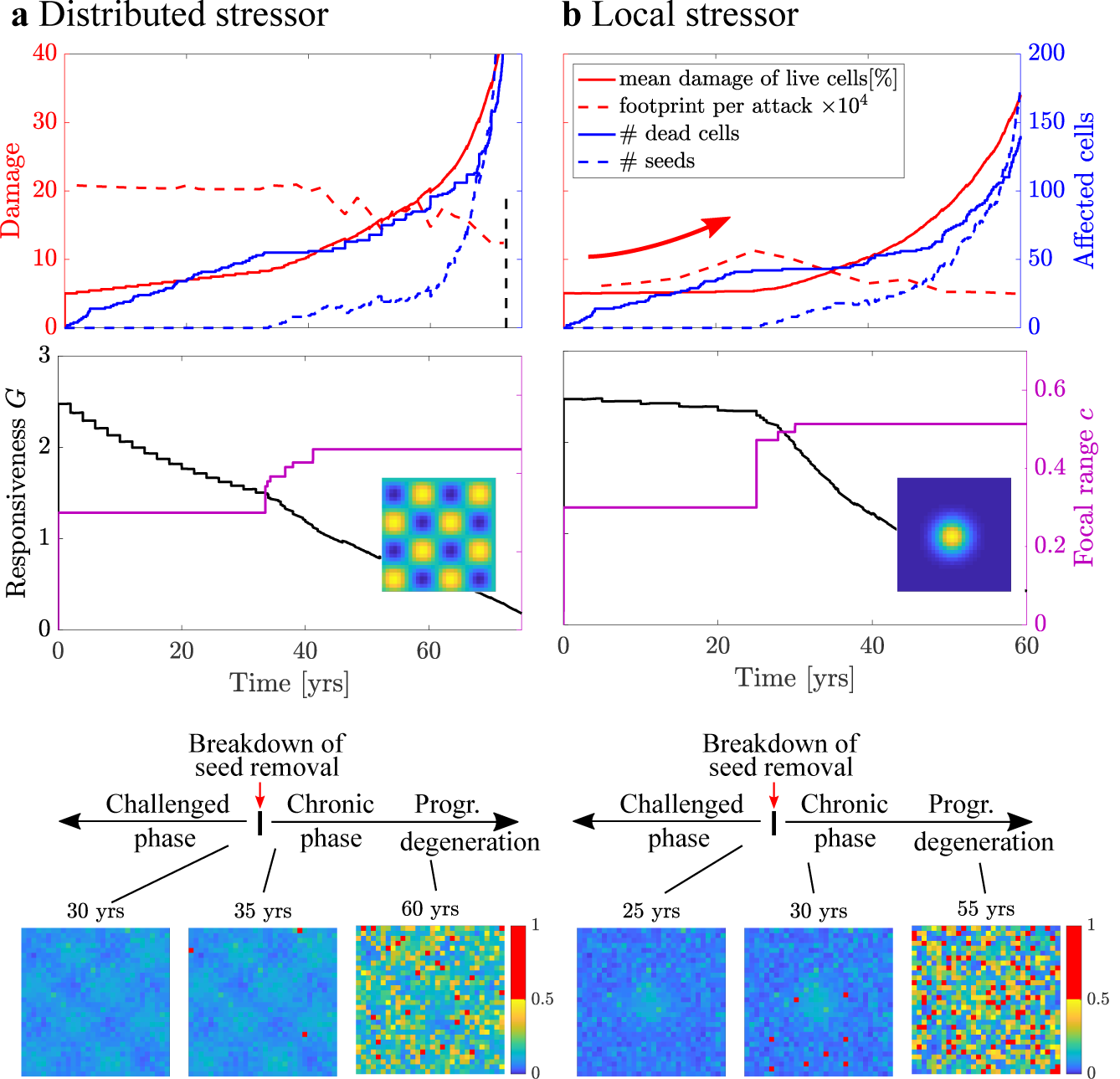
Long-term stress exposure. **Left:** Frequent, weak distributed stressor across the tissue. Gradual adaptation of reactivity *G* to higher mean damage levels leads to a breakdown of seed removal. **Right:** Repeated, stronger stressor at the center of the domain. Repeated local attacks require an adaptation of the focal range *c* to maintain short-term stability, leading to a breakdown of seed removal. Prior to this breakdown, the footprint of each perturbation increases (see red arrows). In both scenarios, after the breakdown of seed removal, cell death is first offset because seeds are maintained in the tissue which leads to increasing damage levels. Eventually, cell death rates increase too. Bottom: Baseline damage levels *z* prior to, immediately after, and long after breakdown of seed removal. Seeds are shown in red. The insets of the graphs of the second row depict the imposed stress pattern. Here, *τ*_∞_ = 50yrs. For model computations without adaptation of *G* and *c*, see Fig. S6. The following randomly induced stressors were induced: (A) Seeds are induced independently with frequency *ν* = 1*/*(10*τ*_∞_). (B) Subthreshold stressors *f*_ext,*n*_(*t*) = *s*(**r**_*n*_) ∑_*k*_ *h*(*t* − *t*_*k*_) are added to the acute cell state are induced at times *t*_*k*_ at rate 1*/*(2yrs) and 1*/*(5yrs). Localized subthreshold stressors are modeled with 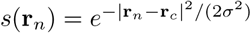 and **r**_*c*_ = (*n/*2, *n/*2). Distributed subthreshold stressors in Fig. S5a are modeled with *s*(**r**_*n*_) = (1 + sin(4*π*(**r**_*n*_)_1_*/n*) sin(4*π*(**r**_*n*_)_2_*/n*))*/*2. The temporal evolution is *h*(*t*) = *Ã*Θ(*t*)(1 + tanh((12h − *t*)*/*4h))*/*2 with heaviside function Θ(*t*) and *Ã*= 0.1. After equilibration after each subthreshold stressor at times *t*_*k*_, baseline damage in increased by 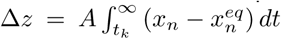, where 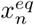 is the equilibrium state before induction of the subthreshold stressor. *A* = 1*/*500 and 1*/*200 for distributed and local stressors, respectively. After each subthreshold stressor, the parameters *c* and *G* were adapted such that: 4*G*⟨*z*⟩ < *H* = 0.5. The focal range was adapted such that the maximal eigenvalue *λ* of Eq. (2) remains *λ* < −0.1*/τ*.

**Figure S6:**
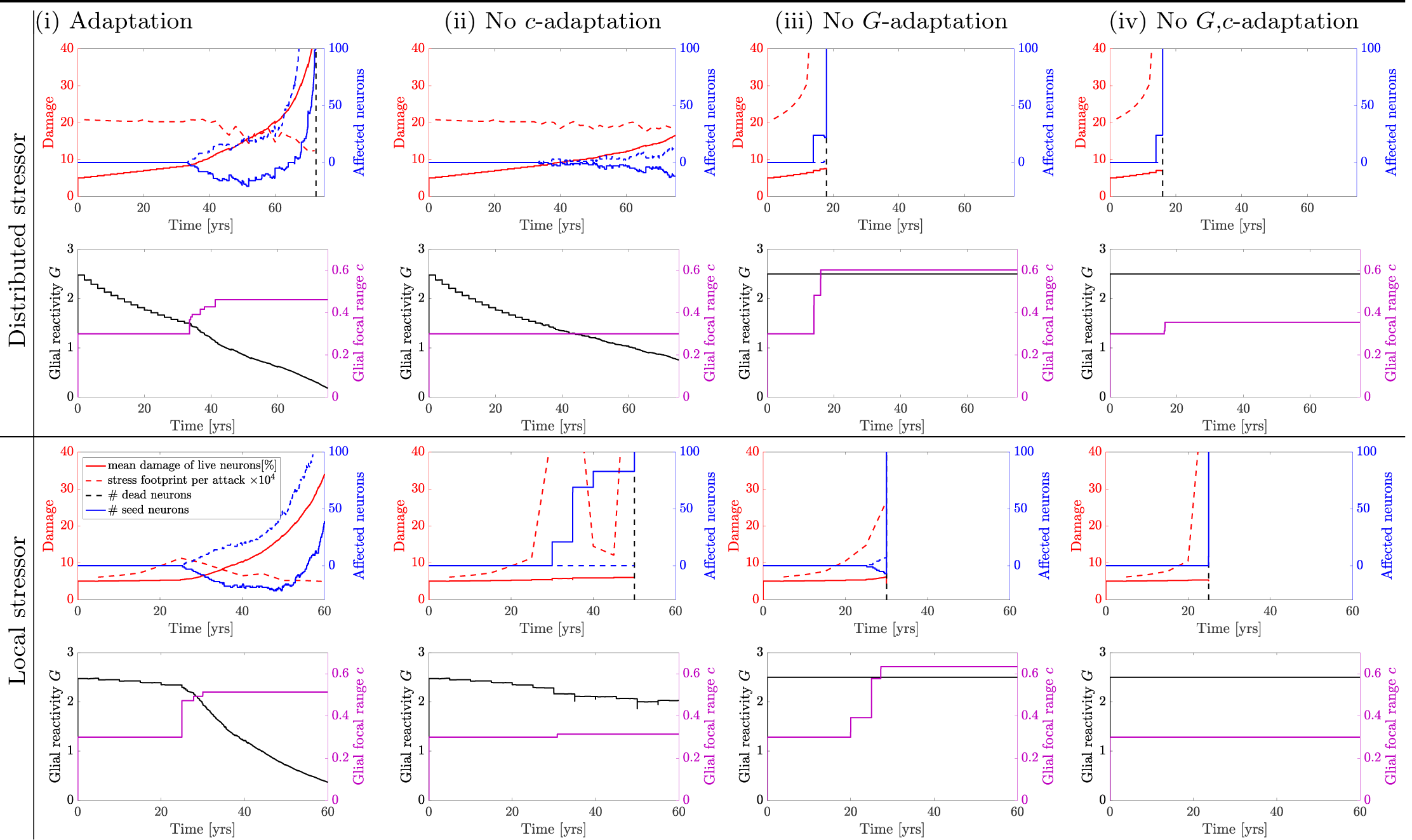
Lack of adaptation leads to early catastrophic cell loss: Effect of adaptation of responsiveness *G* and focal range *c* for distributed and local stressors as in Fig. S5. The first column depicts the results as in Fig. S5 of the main manuscript. In the subsequent columns, we deactivate adaptation of *c* (ii), adaptation of *G* (iii), and both *G* and *c* (iv). In all cases, *c* is not allowed to adapt beyond 0.6. The black dashed line represents the time point of catastrophic cell loss of more than 100 cells (11%). It is seen how no adaptation leads to early catastrophic cell loss. The distributed stressor model is stabilized by *G*-adaptation, and the local stressor model is stabilized by both *G* and *c* adaptation.

## References

[1] Alexey Kolodkin, Evangelos Simeonidis, Rudi Balling, and Hans Westerhoff. Understanding complexity in neurodegenerative diseases: in silico reconstruction of emergence. Frontiers in physiology, 3:291, 2012.

[2] Warren D Anderson and Rajanikanth Vadigepalli. Modeling cytokine regulatory network dynamics driving neuroinflammation in central nervous system disorders. Drug Discovery Today: Disease Models, 19:59–67, 2016.

[3] A Lloret-Villas, TM Varusai, N Juty, C Laibe, N Le Novere, H Hermjakob, and V Chelliah. The impact of mathematical modeling in understanding the mechanisms underlying neurodegeneration: evolving dimensions and future directions. CPT: pharmacometrics & systems pharmacology, 6(2):73–86, 2017.

[4] Mathieu Cloutier, Fiachra B Bolger, John P Lowry, and Peter Wellstead. An integrative dynamic model of brain energy metabolism using in vivo neurochemical measurements. Journal of computational neuroscience, 27(3):391, 2009.

[5] Nathan E Lewis, Gunnar Schramm, Aarash Bordbar, Jan Schellenberger, Michael P Andersen, Jeffrey K Cheng, Nilam Patel, Alex Yee, Randall A Lewis, Roland Eils, et al. Large-scale in silico modeling of metabolic interactions between cell types in the human brain. Nature biotechnology, 28(12):1279, 2010.

[6] Mathieu Cloutier and Peter Wellstead. Dynamic modelling of protein and oxidative metabolisms simulates the pathogenesis of parkinson’s disease. IET systems biology, 6(3):65–72, 2012.

[7] Eleftherios Ouzounoglou, Dimitrios Kalamatianos, Evangelia Emmanouilidou, Maria Xilouri, Leonidas Stefanis, Kostas Vekrellis, and Elias S Manolakos. In silico modeling of the effects of alpha-synuclein oligomerization on dopaminergic neuronal homeostasis. BMC systems biology, 8(1):54, 2014.

[8] Prabha Garg and Jitender Verma. In silico prediction of blood brain barrier permeability: an artificial neural network model. Journal of chemical information and modeling, 46(1):289–297, 2006.

[9] Ines Filipa Martins, Ana L Teixeira, Luis Pinheiro, and Andre O Falcao. A bayesian approach to in silico blood-brain barrier penetration modeling. Journal of chemical information and modeling, 52(6):1686–1697, 2012.

[10] Carole J Proctor and Douglas A Gray. Gsk3 and p53-is there a link in alzheimer’s disease? Molecular neurodegeneration, 5(1):7, 2010.

[11] Theresa M Yuraszeck, Pierre Neveu, Maria Rodriguez-Fernandez, Anne Robinson, Kenneth S Kosik, and Francis J Doyle III. Vulnerabilities in the tau network and the role of ultrasensitive points in tau pathophysiology. PLoS computational biology, 6(11), 2010.

[12] Martin Fussenegger, James E Bailey, and Jeffrey Varner. A mathematical model of caspase function in apoptosis. Nature biotechnology, 18(7):768–774, 2000.

[13] Matthew Y Tang, Carole J Proctor, John Woulfe, and Douglas A Gray. Experimental and computational analysis of polyglutamine-mediated cytotoxicity. PLoS computational biology, 6(9), 2010.

[14] Carole J Proctor and Ian AJ Lorimer. Modelling the role of the hsp70/hsp90 system in the maintenance of protein homeostasis. PloS one, 6(7), 2011.

[15] Gregory Constantine, Marius Buliga, Qi Mi, Florica Constantine, Andrew Abboud, Ruben Zamora, Ava Puccio, David Okonkwo, and Yoram Vodovotz. Dynamic profiling: modeling the dynamics of inflammation and predicting outcomes in traumatic brain injury patients. Frontiers in pharmacology, 7:383, 2016.

[16] Andrew Abboud, Qi Mi, Ava Puccio, David Okonkwo, Marius Buliga, Gregory Constantine, and Yoram Vodovotz. Inflammation following traumatic brain injury in humans: insights from data-driven and mechanistic models into survival and death. Frontiers in pharmacology, 7:342, 2016.

[17] Viktor K Jirsa, William C Stacey, Pascale P Quilichini, Anton I Ivanov, and Christophe Bernard. On the nature of seizure dynamics. Brain, 137(8):2210–2230, 2014.

[18] Anton V Chizhov, Artyom V Zefirov, Dmitry V Amakhin, Elena Yu Smirnova, and Aleksey V Zaitsev. Minimal model of interictal and ictal discharges “epileptor-2”. PLoS computational biology, 14(5):e1006186, 2018.

[19] Honghui Zhang and Pengcheng Xiao. Seizure dynamics of coupled oscillators with epileptor field model. International Journal of Bifurcation and Chaos, 28(03):1850041, 2018.

[20] Kenza El Houssaini, Christophe Bernard, and Viktor K Jirsa. The epileptor model: a systematic mathematical analysis linked to the dynamics of seizures, refractory status epilepticus and depolarization block. Eneuro, 2020.

[21] Epinov. http://www.epinov.com. Accessed: 2020-06-26.

[22] Johannes Weickenmeier, Ellen Kuhl, and Alain Goriely. Multiphysics of prionlike diseases: Progression and atrophy. Physical review letters, 121(15):158101, 2018.

[23] Yasser Iturria-Medina, Roberto C Sotero, Paule J Toussaint, Alan C Evans, Alzheimer’s Disease Neuroimaging Initiative, et al. Epidemic spreading model to characterize misfolded proteins propagation in aging and associated neurodegenerative disorders. PLoS Comput Biol, 10(11):e1003956, 2014.

[24] Jacob W Vogel, Yasser Iturria-Medina, Olof T Strandberg, Ruben Smith, Elizabeth Levitis, Alan C Evans, and Oskar Hansson. Spread of pathological tau proteins through communicating neurons in human alzheimer’s disease. Nature Communications, 11(1):1–15, 2020.

[25] Norihito Uemura, Maiko T Uemura, Kelvin C Luk, Virginia M-Y Lee, and John Q Trojanowski. Cell-to-cell transmission of tau and *α*-synuclein. Trends in Molecular Medicine, 2020.

[26] Guy C. Brown and Jonas J. Neher. Microglial phagocytosis of live neurons. Nat. Rev. Neurosci., 15(4):209–216, 2014.

[27] Knut Biber, Harald Neumann, Kazuhide Inoue, and Hendrikus WGM Boddeke. Neuronal ‘on’ and ‘off’ signals control microglia. Trends Neurosci., 30(11):596–602, 2007.

[28] Richard M. Ransohoff. A polarizing question: do M1 and M2 microglia exist? Nat. Neurosci., 19(8):987–991, 2016.

[29] Nick Stence, Marc Waite, and Michael E. Dailey. Dynamics of microglial activation: a confocal time-lapse analysis in hippocampal slices. Glia, 33:256–266, 2001.

[30] Jan Tønnesen, VVG Krishna Inavalli, and U Valentin Nä gerl. Super-resolution imaging of the extracellular space in living brain tissue. Cell, 172(5):1108–1121, 2018.

[31] John Savill and Valerie Fadok. Corpse clearance defines the meaning of cell death. Nature, 407(6805):784–88, 2000.

[32] Georg W. Kreutzberg. Microglia: a sensor for pathological events in the CNS. Trends Neurosci., 19(8):312–318, 1996.

[33] Yochai Wolf, Simon Yona, Ki-Wook Kim, and Steffen Jung. Microglia, seen from the CX3CR1 angle. Front. Cell. Neurosci., 7:26, 2013.

[34] Helmut Kettenmann, Uwe-Karsten Hanisch, Mami Noda, and Alexei Verkhratsky. Physiology of microglia. Physiol. Rev., 91(2):461–553, 2011.

[35] Jennifer L Zamanian, Lijun Xu, Lynette C Foo, Navid Nouri, Lu Zhou, Rona G Giffard, and Ben A Barres. Genomic analysis of reactive astrogliosis. J. Neurosci., 32(18):6391–6410, 2012.

[36] Baljit S. Khakh and Michael V. Sofroniew. Diversity of astrocyte functions and phenotypes in neural circuits. Nat. Neurosci., 18(7):942–952, 2015.

[37] Michael T Heneka, Róisín M McManus, and Eicke Latz. Inflammasome signalling in brain function and neurodegenerative disease. Nat. Rev. Neurosci., 19(10):610–621, 2018.

[38] Axel Nimmerjahn, Frank Kirchhoff, and Fritjof Helmchen. Resting microglial cells are highly dynamic surveillants of brain parenchyma in vivo. Science, 308:1314–1318, 2005.

[39] Suzanne Hickman, Saef Izzy, Pritha Sen, Liza Morsett, and Joseph El Khoury. Microglia in neurodegeneration. Nat. Neurosci., 21(10):1359, 2018.

[40] Melinda Czeh, Pierre Gressens, and Angela M Kaindl. The yin and yang of microglia. Dev. Neurosci., 33(3-4):199–209, 2011.

[41] G Jean Harry. Microglia during development and aging. Pharm. Ther., 139(3):313–326, 2013.

[42] Michael V Sofroniew and Harry V Vinters. Astrocytes: biology and pathology. Acta Neuropathol., 119(1):7–35, 2010.

[43] Shane A. Liddelow, Kevin A. Guttenplan, Laura E. Clarke, Frederick C. Bennett, Christopher J. Bohlen, Lucas Schirmer, Mariko L. Bennett, Alexandra E. Mönch, Won-Suk Chung, Todd C. Peterson, et al. Neurotoxic reactive astrocytes are induced by activated microglia. Nature, 541(7638):481–487, 2017.

[44] Guy C Brown and Jonas J Neher. Inflammatory neurodegeneration and mechanisms of microglial killing of neurons. Mol. Neurobiol., 41(2-3):242–247, 2010.

[45] Dimitrios Davalos, Jae Kyu Ryu, Mario Merlini, Kim M Baeten, Natacha Le Moan, Mark A Petersen, Thomas J Deerinck, Dimitri S Smirnoff, Catherine Bedard, Hiroyuki Hakozaki, et al. Fibrinogen-induced perivascular microglial clustering is required for the development of axonal damage in neuroinflammation. Nat. Commun., 3:1227, 2012.

[46] Mausam R Damani, Lian Zhao, Aurora M Fontainhas, Juan Amaral, Robert N Fariss, and Wai T Wong. Age-related alterations in the dynamic behavior of microglia. Aging cell, 10(2):263–276, 2011.

[47] Jane H.-C. Lin, Helga Weigel, Maria Luisa Cotrina, Shujun Liu, Earl Bueno, Anker J. Hansen, Thomas W. Hansen, Steven Goldman, and Maiken Nedergaard. Gap-junction-mediated propagation and amplification of cell injury. Nat. Neurosci., 1(6):494–500, 1998.

[48] Taizen Nakase, Shinji Fushiki, and Christian CG Naus. Astrocytic gap junctions composed of connexin 43 reduce apoptotic neuronal damage in cerebral ischemia. Stroke, 34(8):1987–1993, 2003.

[49] Jane H-C Lin, Nanhong Lou, Ning Kang, Takahiro Takano, Furong Hu, Xiaoning Han, Qiwu Xu, Ditte Lovatt, Arnulfo Torres, Klaus Willecke, et al. A central role of connexin 43 in hypoxic preconditioning. J. Neurosci., 28(3):681–695, 2008.

[50] Christian Giaume, Annette Koulakoff, Lisa Roux, David Holcman, and Nathalie Rouach. Astroglial networks: a step further in neuroglial and gliovascular interactions. Nat. Rev. Neurosci., 11(2):87–99, 2010.

[51] Michael R Elliott and Kodi S Ravichandran. The dynamics of apoptotic cell clearance. Developmental cell, 38(2):147–160, 2016.

[52] Susan Elmore. Apoptosis: a review of programmed cell death. Toxicol. Pathol., 35:495–516, 2007.

[53] Brad R. S. Broughton, David C. Reutens, and Christopher G. Sobey. Apoptotic mechanisms after cerebral ischemia. Stroke, 40(5):e331–e339, 2009.

[54] Mark A Petersen and Michael E Dailey. Diverse microglial motility behaviors during clearance of dead cells in hippocampal slices. Glia, 46(2):195–206, 2004.

[55] Anke Witting, Peter Möller, Andreas Herrmann, Helmut Kettenmann, and Christiane Nolte. Phagocytic clearance of apoptotic neurons by microglia/brain macrophages in vitro. Journal of neurochemistry, 75(3):1060–1070, 2000.

[56] Sanja Arandjelovic and Kodi S Ravichandran. Phagocytosis of apoptotic cells in homeostasis. Nature immunology, 16(9):907–917, 2015.

[57] Pinar Ayata, Ana Badimon, Hayley J Strasburger, Mary Kaye Duff, Sarah E Montgomery, Yong-Hwee E Loh, Anja Ebert, Anna A Pimenova, Brianna R Ramirez, Andrew T Chan, et al. Epigenetic regulation of brain region-specific microglia clearance activity. Nat. Neurosci., 21(8):1049, 2018.

[58] Ivan K. H. Poon, Christopher D. Lucas, Adriano G. Rossi, and Kodi S. Ravichandran. Apoptotic cell clearance: basic biology and therapeutic potential. Nat. Rev. Immunol., 14(3):166–180, 2014.

[59] H. Neumann, M. R. Kotter, and R. J. M. Franklin. Debris clearance by microglia: an essential link between degeneration and regeneration. Brain, 132(2):288–295, 2008.

[60] I. Napoli and H. Neumann. Microglial clearance function in health and disease. Neuroscience, 158(3):1030–1038, 2009.

[61] Francesco Di Virgilio, Stefania Ceruti, Placido Bramanti, and Maria P. Abbracchio. Purinergic signalling in inflammation of the central nervous system. Trends Neurosci., 32(2):79–87, 2009.

[62] Dimitry Ofengeim, Yasushi Ito, Ayaz Najafov, Yaoyang Zhang, Bing Shan, Judy Park DeWitt, Juanying Ye, Xumin Zhang, Ansi Chang, Helin Vakifahmetoglu-Norberg, et al. Activation of necroptosis in multiple sclerosis. Cell reports, 10(11):1836–1849, 2015.

[63] Antonella Caccamo, Caterina Branca, Ignazio S Piras, Eric Ferreira, Matthew J Huentelman, Winnie S Liang, Ben Readhead, Joel T Dudley, Elizabeth E Spangenberg, Kim N Green, et al. Necroptosis activation in alzheimer’s disease. Nature Neuroscience, 20(9):1236–1246, 2017.

[64] Li Gan, Mark R Cookson, Leonard Petrucelli, and Albert R La Spada. Converging pathways in neurodegeneration, from genetics to mechanisms. Nat. Neurosci., 21(10):1300, 2018.

[65] Don H Mahad, Bruce D Trapp, and Hans Lassmann. Pathological mechanisms in progressive multiple sclerosis. Lancet Neurol., 14(2):183–193, 2015.

[66] Yongdae Shin and Clifford P Brangwynne. Liquid phase condensation in cell physiology and disease. Science, 357(6357):eaaf4382, 2017.

[67] Hongjun Fu, John Hardy, and Karen E Duff. Selective vulnerability in neurodegenerative diseases. Nat. Neurosci., 21(10):1350–1358, 2018.

[68] Carmen Venegas, Sathish Kumar, Bernardo S. Franklin, Tobias Dierkes, Rebecca Brinkschulte, Dario Tejera, Ana Vieira-Saecker, Stephanie Schwartz, Francesco Santarelli, Markus P. Kummer, et al. Microglia-derived ASC specks cross-seed amyloid-*β* in Alzheimer’s disease. Nature, 552(7685):355–361, 2017.

[69] Mathias Jucker and Lary C Walker. Propagation and spread of pathogenic protein assemblies in neurodegenerative diseases. Nat. Neurosci., 21(10):1341, 2018.

[70] Michelle L Block, Luigi Zecca, and Jau-Shyong Hong. Microglia-mediated neurotoxicity: uncovering the molecular mechanisms. Nat. Rev. Neurosci., 8(1):57, 2007.

[71] Junying Yuan, Palak Amin, and Dimitry Ofengeim. Necroptosis and RIPK1-mediated neuroinflammation in CNS diseases. Nat. Rev. Neurosci., 20(1):19–33, 2018.

[72] Thomas F Gajewski, Hans Schreiber, and Yang-Xin Fu. Innate and adaptive immune cells in the tumor microenvironment. Nature immunology, 14(10):1014–1022, 2013.

[73] Peng Jiang, Shengqing Gu, Deng Pan, Jingxin Fu, Avinash Sahu, Xihao Hu, Ziyi Li, Nicole Traugh, Xia Bu, Bo Li, et al. Signatures of t cell dysfunction and exclusion predict cancer immunotherapy response. Nature medicine, 24(10):1550–1558, 2018.

[74] Christopher M Henstridge, Bradley T Hyman, and Tara L Spires-Jones. Beyond the neuron–cellular interactions early in Alzheimer disease pathogenesis. Nat. Rev. Neurosci., 20(2):94–108, 2019.

[75] Celeste M Karch and Alison M Goate. Alzheimer’s disease risk genes and mechanisms of disease pathogenesis. Biol. Psychiatry, 77(1):43–51, 2015.

[76] Cláudia Y. Santos, Peter J. Snyder, Wen-Chih Wu, Mia Zhang, Ana Echeverria, and Jessica Alber. Pathophysiologic relationship between Alzheimer’s disease, cerebrovascular disease, and cardiovascular risk: a review and synthesis. Alzheimers Dement. (AMST), 7:69–87, 2017.

[77] Sandro Da Mesquita, Antoine Louveau, Andrea Vaccari, Igor Smirnov, R. Chase Cornelison, Kathryn M. Kingsmore, Christian Contarino, Suna Onengut-Gumuscu, Emily Farber, Daniel Raper, et al. Functional aspects of meningeal lymphatics in ageing and Alzheimer’s disease. Nature, 560(7717):185–191, 2018.

[78] Michelle L. Block and Jau-Shyong Hong. Microglia and inflammation-mediated neurodegeneration: multiple triggers with a common mechanism. Prog. Neurobiol., 76(2):77–98, 2005.

[79] Annett Halle, Veit Hornung, Gabor C. Petzold, Cameron R. Stewart, Brian G. Monks, Thomas Reinheckel, Katherine A. Fitzgerald, Eicke Latz, Kathryn J. Moore, and Douglas T. Golenbock. The NALP3 inflammasome is involved in the innate immune response to amyloid-*β*. Nat. Immunol., 9(8):857–865, 2008.

[80] Frederick J. Sheedy, Alena Grebe, Katey J. Rayner, Parisa Kalantari, Bhama Ramkhelawon, Susan B. Carpenter, Christine E. Becker, Hasini N. Ediriweera, Adam E. Mullick, Douglas T. Golenbock, et al. CD36 coordinates NLRP3 inflammasome activation by facilitating intracellular nucleation of soluble ligands into particulate ligands in sterile inflammation. Nat. Immunol., 14(8):812–820, 2013.

[81] Frank L. Heppner, Richard M. Ransohoff, and Burkhard Becher. Immune attack: the role of inflammation in Alzheimer’s disease. Nat. Rev. Neurosci., 16(6):358–372, 2015.

[82] Michael T Heneka, Monica J Carson, Joseph El Khoury, Gary E Landreth, Frederic Brosseron, Douglas L Feinstein, Andreas H Jacobs, Tony Wyss-Coray, Javier Vitorica, Richard M Ransohoff, et al. Neuroinflammation in Alzheimer’s disease. Lancet Neurol., 14(4):388–405, 2015.

[83] Michael T Heneka, Douglas T Golenbock, and Eicke Latz. Innate immunity in Alzheimer’s disease. Nature Immunol., 16(3):229, 2015.

[84] Maike Gold and Joseph El Khoury. *β*-amyloid, microglia, and the inflammasome in Alzheimer’s disease. In Semin. Immunopathol., volume 37, pages 607–611. Springer, 2015.

[85] Joseph B El Khoury, Kathryn J Moore, Terry K Means, Josephine Leung, Kinya Terada, Michelle Toft, Mason W Freeman, and Andrew D Luster. CD36 mediates the innate host response to *β*-amyloid. J. Exp. Med., 197(12):1657–1666, 2003.

[86] Indra Sethy Coraci, Jens Husemann, Joan W Berman, Christine Hulette, Jennifer H Dufour, Gabriele K Campanella, Andrew D Luster, Samuel C Silverstein, and Joseph B El Khoury. CD36, a class B scavenger receptor, is expressed on microglia in Alzheimer’s disease brains and can mediate production of reactive oxygen species in response to *β*-amyloid fibrils. Am. J. Pathol., 160(1):101–112, 2002.

[87] Rommy Von Bernhardi, Laura Eugenín-von Bernhardi, and Jaime Eugenín. Microglial cell dysregulation in brain aging and neurodegeneration. Front. Aging Neurosci., 7:124, 2015.

[88] Elizabeth E. Spangenberg, Rafael J. Lee, Allison R. Najafi, Rachel A. Rice, Monica R. P. Elmore, Mathew Blurton-Jones, Brian L. West, and Kim N. Green. Eliminating microglia in Alzheimer’s mice prevents neuronal loss without modulating amyloid-*β* pathology. Brain, 139:1265–1281, 2016.

[89] Bart De Strooper and Eric Karran. The cellular phase of Alzheimer’s disease. Cell, 164(4):603–615, 2016.

[90] Rommy Von Bernhardi. Glial cell dysregulation: a new perspective on Alzheimer disease. Neurotox Res., 12(4):215–232, 2007.

[91] Alix De Calignon, Manuela Polydoro, Marc Suárez-Calvet, Christopher William, David H Adamowicz, Kathy J Kopeikina, Rose Pitstick, Naruhiko Sahara, Karen H Ashe, George A Carlson, et al. Propagation of tau pathology in a model of early Alzheimer’s disease. Neuron, 73(4):685–697, 2012.

[92] Johannes Brettschneider, Kelly Del Tredici, Virginia M.-Y. Lee, and John Q. Trojanowski. Spreading of pathology in neurodegenerative diseases: a focus on human studies. Nat. Rev. Neurosci., 16(2):109–120, 2015.

[93] D. James Surmeier, José A. Obeso, and Glenda M. Halliday. Selective neuronal vulnerability in Parkinson disease. Nat. Rev. Neurosci., 18(2):101–113, 2017.

[94] Jing L Guo and Virginia MY Lee. Cell-to-cell transmission of pathogenic proteins in neurodegenerative diseases. Nat. Med., 20(2):130, 2014.

[95] Marc Aurel Busche, Gerhard Eichhoff, Helmuth Adelsberger, Dorothee Abramowski, Karl-Heinz Wiederhold, Christian Haass, Matthias Staufenbiel, Arthur Konnerth, and Olga Garaschuk. Clusters of hyperactive neurons near amyloid plaques in a mouse model of alzheimer’s disease. Science, 321(5896):1686–1689, 2008.

[96] Benedikt Zott, Manuel M Simon, Wei Hong, Felix Unger, Hsing-Jung Chen-Engerer, Matthew P Frosch, Bert Sakmann, Dominic M Walsh, and Arthur Konnerth. A vicious cycle of *β* amyloid–dependent neuronal hyperactivation. Science, 365(6453):559–565, 2019.

[97] Samira Parhizkar, Thomas Arzberger, Matthias Brendel, Gernot Kleinberger, Maximilian Deussing, Carola Focke, Brigitte Nuscher, Monica Xiong, Alireza Ghasemigharagoz, Natalie Katzmarski, et al. Loss of TREM2 function increases amyloid seeding but reduces plaque-associated apoe. Nat. Neurosci., 22(2):191–204, 2019.

[98] Wilbur M Song, Satoru Joshita, Yingyue Zhou, Tyler K Ulland, Susan Gilfillan, and Marco Colonna. Humanized TREM2 mice reveal microglia-intrinsic and-extrinsic effects of R47H polymorphism. J. Exp. Med., 215(3):745–760, 2018.

[99] Fred D. Lublin, Stephen C. Reingold, Jeffrey A. Cohen, Gary R. Cutter, Per Soelberg Sørensen, Alan J. Thompson, Jerry S. Wolinsky, Laura J. Balcer, Brenda Banwell, Frederik Barkhof, et al. Defining the clinical course of multiple sclerosis. The 2013 revisions. Neurology, 83(3):278–286, 2014.

[100] Volker Siffrin, Helena Radbruch, Robert Glumm, Raluca Niesner, Magdalena Paterka, Josephine Herz, Tina Leuenberger, Sabrina M. Lehmann, Sarah Luenstedt, Jan Leo Rinnenthal, et al. In vivo imaging of partially reversible Th17 cell-induced neuronal dysfunction in the course of encephalomyelitis. Immunity, 33(3):424–436, 2010.

[101] Roland S. Liblau, Daniel Gonzalez-Dunia, Heinz Wiendl, and Frauke Zipp. Neurons as targets for T cells in the nervous system. Trends Neurosci., 36(6):315–324, 2013.

[102] Catherine Larochelle, Timo Uphaus, Alexandre Prat, and Frauke Zipp. Secondary progression in multiple sclerosis: neuronal exhaustion or distinct pathology? Trends Neurosci., 39(5):325–339, 2016.

[103] Thomas Korn, Tim Magnus, and Stefan Jung. Autoantigen specific T cells inhibit glutamate uptake in astrocytes by decreasing expression of astrocytic glutamate transporter GLAST: a mechanism mediated by tumor necrosis factor-*α*. The FASEB journal, 19(13):1878–1880, 2005.

[104] K. Mitosek-Szewczyk, G. Sulkowski, Z. Stelmasiak, and L. Strużyńska. Expression of glutamate transporters GLT-1 and GLAST in different regions of rat brain during the course of experimental autoimmune encephalomyelitis. Neuroscience, 155(1):45–52, 2008.

[105] Nicholas J. Maragakis and Jeffrey D. Rothstein. Mechanisms of disease: astrocytes in neurodegenerative disease. Nat. Rev. Neurol., 2(12):679–689, 2006.

[106] Geraldine T Petr, Yan Sun, Natalie M Frederick, Yun Zhou, Sameer C Dhamne, Mustafa Q Hameed, Clive Miranda, Edward A Bedoya, Kathryn D Fischer, Wencke Armsen, et al. Conditional deletion of the glutamate transporter GLT-1 reveals that astrocytic GLT-1 protects against fatal epilepsy while neuronal GLT-1 contributes significantly to glutamate uptake into synaptosomes. J. Neurosci., 35(13):5187–5201, 2015.

[107] Dirk Luchtman, René Gollan, Erik Ellwardt, Jérôme Birkenstock, Kerstin Robohm, Volker Siffrin, and Frauke Zipp. In vivo and in vitro effects of multiple sclerosis immunomodulatory therapeutics on glutamatergic excitotoxicity. J. Neurochem., 136:971–980, 2016.

[108] Khalil S Rawji, Janson Kappen, Weiwen Tang, Wulin Teo, Jason R Plemel, Peter K Stys, and V Wee Yong. Deficient surveillance and phagocytic activity of myeloid cells within demyelinated lesions in ageing mice visualized by ex vivo live multiphoton imaging. J. Neurosci., pages 2341–17, 2018.

[109] Ludovico Cantuti-Castelvetri, Dirk Fitzner, Mar Bosch-Queralt, Marie-Theres Weil, Minhui Su, Paromita Sen, Torben Ruhwedel, Miso Mitkovski, George Trendelenburg, Dieter Lö tjohann, et al. Defective cholesterol clearance limits remyelination in the aged central nervous system. Science, 359(6376):684–688, 2018.

[110] Robin JM Franklin et al. Regenerating CNS myelin: from mechanisms to experimental medicines. Nat. Rev. Neurosci., 18(12):753, 2017.

[111] Mohamed El Behi, Charles Sanson, Corinne Bachelin, Léna Guillot-Noë l, Jennifer Fransson, Bruno Stankoff, Elisabeth Maillart, Nadège Sarrazin, Vincent Guillemot, Hervé Abdi, et al. Adaptive human immunity drives remyelination in a mouse model of demyelination. Brain, 140(4):967–980, 2017.

[112] Bernhard Hemmer, Martin Kerschensteiner, and Thomas Korn. Role of the innate and adaptive immune responses in the course of multiple sclerosis. Lancet Neurol., 14(4):406–419, 2015.

[113] Klaus-Armin Nave. Myelination and support of axonal integrity by glia. Nature, 468(7321):244–252, 2010.

[114] Benjamin Schattling, Jan Broder Engler, Constantin Volkmann, Nicola Rothammer, Marcel S Woo, Meike Petersen, Iris Winkler, Max Kaufmann, Sina C Rosenkranz, Anna Fejtova, et al. Bassoon proteinopathy drives neurodegeneration in multiple sclerosis. Nat. Neurosci., page 1, 2019.

[115] Mohammad Rohani and Shadi Ghourchian. Fulminant multiple sclerosis (MS). Neurol. Sci., 32(953), 2011.

[116] Philip L De Jager, Hyun-Sik Yang, and David A Bennett. Deconstructing and targeting the genomic architecture of human neurodegeneration. Nat. Neurosci., 21(10):1310, 2018.

[117] Aleksandra Deczkowska, Ido Amit, and Michal Schwartz. Microglial immune checkpoint mechanisms. Nat. Neurosci., 21(6):779–786, 2018.

[118] Raffaella Nativio, Greg Donahue, Amit Berson, Yemin Lan, Alexandre Amlie-Wolf, Ferit Tuzer, Jon B. Toledo, Sager J. Gosai, Brian D. Gregory, Claudio Torres, et al. Dysregulation of the epigenetic landscape of normal aging in Alzheimer’s disease. Nat. Neurosci., 21(4):497–505, 2018.

[119] Marten Scheffer, Stephen R. Carpenter, Timothy M. Lenton, Jordi Bascompte, William Brock, Vasilis Dakos, Johan van de Koppel, Ingrid A. van de Leemput, Simon A. Levin, Egbert H. van Nes, et al. Anticipating critical transitions. Science, 338:344–348, 2012.

[120] Renaud Jolivet, Jay S Coggan, Igor Allaman, and Pierre J Magistretti. Multi-timescale modeling of activity-dependent metabolic coupling in the neuron-glia-vasculature ensemble. PLoS Comput Biol, 11(2):e1004036, 2015.

[121] Mathias Jucker and Lary C Walker. Self-propagation of pathogenic protein aggregates in neurodegenerative diseases. Nature, 501(7465):45, 2013.

[122] Johannes Weickenmeier, Ellen Kuhl, and Alain Goriely. Multiphysics of prionlike diseases: Progression and atrophy. Phys. Rev. Lett., 121:158101, Oct 2018.

